# Introgression and divergence in a young species group

**DOI:** 10.1101/2024.12.23.630027

**Authors:** I. Satokangas, SH. Martin, B. Seifert, T. Puukko, H. Helanterä, J. Kulmuni

**Affiliations:** Organismal & Evolutionary Biology Research Programme, University of Helsinki, Viikinkaari 1, P.O.Box 65, 00014 University of Helsinki, Finland; Institute of Evolutionary Biology, University of Edinburgh, Ashworth Laboratories, Edinburgh EH9 3FL, UK; Department of Entomology, Senckenberg Museum für Naturkunde, Am Museum 1, D-02826 Görlitz, Germany; Ecology and Genetics research unit, University of Oulu, P.O. Box 3000, 90014 University of Oulu, Finland; Tvärminne Zoological Station, University of Helsinki, J.A. Palménin tie 260, FI-10900 Hanko, Finland; Institute for Biodiversity and Ecosystem Dynamics, Department of Evolutionary and Population Biology, University of Amsterdam, Amsterdam, The Netherlands

**Keywords:** Speciation, hybridization, introgression, divergence, *Formica* wood ants

## Abstract

The process of speciation concerns often not only pairs of species but rather groups of diverging and interacting taxa, as highlighted by recent research. Hence, to understand the evolution of species’ diversity and their persistence, it is crucial to understand how gene flow and evolution of reproductive isolation shape groups of closely related species. Using resequencing data, we disentangle here genomic patterns of divergence and introgression in five *Formica rufa* group wood ant species that are at the early stage of speciation. We first revise earlier mitochondrial phylogenies with a nuclear genomic tree, and demonstrate then introgression that is in line with observations of their current day natural hybridisation. Investigating the genome-wide differentiation and divergence we find correlations between population genetic parameters of divergence, differentiation, and diversity, that are in line with theoretical expectations for young species. Despite previously found evidence for polygenic species barriers, our data lacks the genome-wide correlation between differentiation and divergence that would be expected under a model of polygenic barriers. The likely explanation for this lack is the dominating effect of ancestral diversity at these early stages of speciation. As hybridisation has led to both deleterious and adaptive consequences within the group, we examined the signatures of introgression. We find no strong positive correlation between introgression and recombination, suggesting introgression does not have a predominantly deleterious effect. We also infer low diversity in the genomic regions with high proportions of introgression, consistent with the idea that selection has locally favoured introgression. This could be due to sharing of adaptive alleles or reduction of genetic load in the receiving species. Interestingly, gene flow in this group could potentially cross multiple species boundaries even in the absence of direct interbreeding between all the species. We discuss the long-term benefits and costs of introgression in young species, including the effect of environmental fluctuations and multi-species introgression.

## 1. Introduction

Recent studies have highlighted that speciation is a reticulate process that involves and is influenced by hybridisation and gene flow. Gene flow, the transfer of genetic material between diverging lineages, may be neutral, beneficial, or deleterious, and either facilitate or slow down divergence and adaptation of incipient species (Peñalba et al. 2024; Abbott et al. 2013). Hybridization and repeated backcrossing may eventually lead to introgression, whereby genetic material from one species is transferred into and modifies the gene pool of another species. It is central to understand the signatures that beneficial and deleterious introgression, or barriers that impede them, leave in the genome during divergence. This helps to identify the genomic regions that are important for adaptation and speciation. A range of empirical examples show how gene flow and introgression may occur between multiple diverging lineages (Suvorov et al. 2022; Meier et al. 2023; Kozak et al. 2021) transferring genetic material even between species which do not directly interbreed with each other (Grant and Grant 2020). Despite this evidence, it is not well known how introgression influences speciation, especially in groups of young taxa.

Currently, it remains unclear how the fitness effects of introgression change over divergence. Conventionally, introgression has been seen as largely deleterious, yet there is a growing body of evidence for introgression of adaptive alleles in various organisms (Oziolor et al. 2019; Norris et al. 2015; Racimo et al. 2015; Pardo-Diaz et al. 2012). According to the conventional view, introgression is most prevalent in regions of high recombination, where small adaptive or neutral parts can be recombined out of mostly deleterious material and hence maintained in the population. This scenario holds according to both theoretical as well as empirical (Sankararaman et al. 2014; Martin et al. 2019) research especially for divergent species.

Only recent theoretical work has addressed the interplay of divergence and introgression, especially how selection towards different sized introgressed haplotypes changes with divergence. This work suggests that early in divergence the fitness effect or large introgressing blocks may be neutral or beneficial instead of being largely deleterious, which would facilitate the introgression of large haplotypes. In young species, the benefits of coadapted sets of alleles within introgressing blocks are suggested to outweigh the disadvantages of weak deleterious epistatic effects with the recipient genome, resulting in novel expectations about the genomic signatures of introgression (Dagilis and Matute 2023). Introgression would be independent of the amount of recombination, or become more prevalent in regions of low recombination where the introgressed blocks can be selected for as they are. Such a pattern has been demonstrated empirically at least in secondary contact (Duranton and Pool 2022; Pool 2015; Dagilis and Matute 2023). If divergence would happen with continuous gene flow, the species would likely become and stay genomically distinct only due to adaptive divergence (see, e.g., Malinsky et al. (2015)), whereby introgression of diverged haplotypes would likely be maladaptive.

Furthermore, an alternative explanation for a negative correlation between recombination and introgression has been suggested in theoretical work modelling the impact of genetic load for introgression. In these models, a negative correlation is not linked to early divergence, but may arise if introgression masks (or purges) accumulated recessive deleterious variation in species with high genetic load (Kim, Huber, and Lohmueller 2018). In this work, we use the term adaptive introgression both when introgression leads to exchange of genetic material that underlies differential adaptation, as well as when introgression is selected for because it helps masking or purging deleterious variation from genetic load. A signature of adaptive introgression is in both cases, at simplest, reduced nucleotide diversity (π) at regions of high introgression. This signals that selection has favoured introgression as demonstrated, for instance, in Arabidopsis (Arnold et al. 2016).

In order to form a thorough understanding of the studied taxa and the signatures different evolutionary processes leave in the genome during divergence, summary statistics of genetic diversity, divergence, and differentiation can be used. In this work, we measure genetic diversity as nucleotide diversity, π, the average number of nucleotide differences per site within a species. By genetic divergence we mean d_xy_, the average number of nucleotide differences per site between species (Nei and Li 1979). Differentiation, on the other hand, refers to *F*_ST_, which measures how similar or different allele frequencies are between two species (differentiation between species relative to the total genetic diversity across both species) (Weir and Cockerham 1984). These measures are expected to develop different correlations depending on the evolutionary scenario and the divergence between the species. In young species, sequence divergence (d_xy_) largely reflects the ancestral diversity (i.e., existing diversity (π) in the ancestral population before the incipient species became separated). The ancestral diversity varies heterogeneously along the genome due to background selection and local mutation rates (Charlesworth, Morgan, and Charlesworth 1993). As both diversity and divergence are computed from pairwise sequence differences, low within-species diversity translates into low between-species divergence, and high diversity into high divergence, creating a strong positive whole-genome correlation that serves as a hallmark of recently split species. Through time, the correlation weakens as both lineages start experiencing increasingly different selection pressures and the patterns of diversity diverge. In the face of strong gene flow, even a negative correlation between diversity and divergence may emerge, as divergence accumulates in genomic regions where divergent selection limits the effective migration rate (Shang ^e^t al. 2023). The relationship between genetic diversity and differentiation (*F*_ST_), on the other hand, may be stochastic in early divergence as differentiation first reflects sampling effect and is created in incipient species due to differences in sorting of the ancestral variation. After this, both positive and background selection are expected to lead to a negative genome-wide correlation between differentiation and diversity that strengthens with increasing divergence (Burri 2017; Stankowski et al. 2019).

As divergence progresses, the prevalence of regions that resist introgression increases (Orr and Turelli 2001; Orr 1995), opposite to the amount of introgression (Hamlin, Hibbins, and Moyle 2020). These so-called gene flow barriers are responsible for speciation when divergence occurs with gene flow. However, how their genomic distribution develops, or whether they are formed of a few regions or scattered along the whole genome has continued to be under investigation for the past decades (Laetsch et al. 2023; Wu 2001). A common way to look for such regions has been to perform so-called outlier scans to detect highly differentiated regions that supposedly arise because they resist gene flow. Such scans measuring only differentiation (*F*_ST_) were originally used to detect the barriers but high differentiation may in fact arise also due to processes unrelated to reproductive isolation (Wolf and Ellegren 2017). Instead, gene flow barriers should manifest as regions where both differentiation and absolute divergence (d_xy_) are elevated - a correlation that is absent or negative in the absence of gene flow (Shang et al. 2023).

We study introgression and divergence using *Formica rufa* group wood ants that have diverged recently within the last ∼500.000 years (Portinha et al. 2022; Goropashnaya, Fedorov, and Pamilo 2004). They live in boreal forests throughout Eurasia with large sympatric areas (Stockan and Robinson 2016). The *F. rufa* group has at least two distinguishable features in which the species differ from each other and that likely contribute to their speciation. First, the non-sister species have different climatic adaptations that are reflected in their geographical distributions (Stockan and Robinson 2016). Second, sister species harbour different social strategies (polygyny and monogyny) that are coupled with their dispersal patterns and habitat requirements (Seifert 2018; Stockan and Robinson 2016). At least two non-sister species have, according to demographic modelling, diverged with continuous gene flow until recent times (Portinha et al. 2022). However, as the *F. rufa* group species have been suggested to diverge for periods of time in separated forest regions, the Pleistocene glacial refugia (Goropashnaya, Fedorov, and Pamilo 2004), their history likely encompasses both periods of allopatry and gene flow. Currently multiple species in the *F. rufa* group hybridise extensively, which has led to formation of further generation and backcrossed hybrid populations (Satokangas et al. 2023; Seifert 2021). Hybridisation of the two non-sister species, *F. aquilonia* and *F. polyctena*, has been coupled with various consequences. These include hybrid mortality (Kulmuni and Pamilo 2014), repeatable sorting of ancestry presumably due to both genetic load and favouring of adaptive alleles (Nouhaud et al. 2022), and potential to adapt to climate change due to differential climatic adaptations (Satokangas et al. 2023; Martin-Roy et al. 2021). However, the overall fitness effects and long-term impacts of introgression are not known, or how speciation and polygenic gene flow barriers detected in previous work (Kulmuni and Pamilo 2014; Kulmuni et al. 2020) manifest as whole-genome divergence in this group.

Using a comparative population genomic framework, we utilise new and pre-existing resequencing data from five wood ant species to understand how different processes have contributed to their speciation during the recent divergence history. We demonstrate that genetic divergence resembles ancestral diversity, signalling recent speciation. We revise earlier mitochondrial phylogenies with a nuclear species tree. Despite a high proportion of unsorted variation, the tree clusters individuals by species regardless of geographic origin. We find that genome-wide correlations between population genetic parameters are in line with theoretical expectations for young species: in addition to strong significant positive correlation between divergence and within-species diversity, there is no signal of barriers to gene flow that would be detectable as a correlation between differentiation and divergence at the whole-genome scale. We detect introgression between sister and non-sister species pairs. Hybridisation in both pairs is extensive in current-day natural populations. In previous work, recent-day hybrids have been shown to suffer from mortality, but they are also suggested to benefit from decreased genetic load and increased potential to adapt to changing climate. Our results on the introgression landscape suggest that the overall fitness effect of large introgressing blocks in these wood ant species is not deleterious as conventionally considered. Inferred introgression is largely neutral and even beneficial, following the expectations for young species from new theoretical work. Neutrality is inferred from at most a very weak positive whole-genome correlation between recombination and introgression. Low diversity in the regions of highest introgression suggest a proportion of it is adaptive, either because introgression masks recessive genetic load, or because of other adaptive value.

## 2. Materials and Methods

### 2.1. Sampling

We studied genomic signatures of diversity, differentiation, divergence, and introgression among five *Formica rufa* group wood ant species. Our aim was to investigate if we find signatures of either adaptive or deleterious introgression among the species group. We also studied the relationship between diversity, divergence and differentiation to produce a thorough understanding of speciation in this group, and constructed an improved whole-genome species tree.

For these purposes, we sampled individual wood ant workers from five sympatrically living *Formica rufa* group species: *F. aquilonia*, *F. lugubris*, *F. polyctena*, *F. rufa*, and *F. pratensis* (Satokangas et al. 2023; Stockan and Robinson 2016). Major part of the samples originate from Finland, supplemented with samples from Central Europe and Scotland to offer a broader view. We utilised whole-genome sequencing data, mostly for one individual per population.

Majority of our data consist of individuals sequenced in previous work (Satokangas et al. 2023; Portinha et al. 2022). In addition to *F. rufa* group samples, *Formica exsecta*, a distantly related species belonging to the *Formica exsecta* group (Borowiec, Cover, and Rabeling 2021), was used as an outgroup in some analyses ((Nouhaud et al. 2022); originally published in Dhaygude et al. (2019)). New *F. rufa* group ant samples with no published genomic data were obtained from Germany and Switzerland, primarily to clarify the relationship of *F. polyctena* and *F. rufa*. These samples were stored in ethanol prior to DNA extraction and classified morphologically. This classification was based on investigation of dry, mounted worker specimens by high-resolution stereomicroscopy considering 16 characters. These characters included one indicator of absolute size, four shape variables, seta counts in six body parts and seta length measurements in five body parts. The samples were classified using the principal component analysis (PCA), and in order to improve the PCA performance, removal of allometric variance was performed for all shape and seta characters for the assumption of each individual having a cephalic size of 1750 µm (for details see Seifert (2021)). See S1 Table for more information on the sampling, including sampling locations and years. Taken together, in this study we analysed altogether 93 *F. rufa* group individuals. Majority of the analyses were performed with five individuals per species, while the remaining individuals served as context for our findings.

### 2.2. Whole-genome sequencing

For the individuals used in previous work, the details of whole-genome sequencing and first data processing steps can be found in Satokangas et al. (2023). For the new samples, we extracted DNA from the whole worker ant bodies. Tissue samples (individual workers) were stored in ethanol and before genomic DNA extraction we removed the ethanol by gently pressing with a paper towel and then we ground the samples in micro centrifuge tubes using liquid nitrogen and plastic pestles. We then extracted DNA with Qiagen’s DNeasy Blood & Tissue Kit (Cat. No. / ID: 69504). We extended the lysis step overnight and did not do the RNase treatment. We quantified DNA concentrations with a ThermoFisher Scientific Qubit DS DNA Kit (Q32851).

We randomised the sample order and then used New England Biolabs NEBNext® Ultra™ II FS DNA Library Prep Kit for Illumina (E7805L) to prepare the DNA libraries, and indexed the samples using NEBNext® Multiplex Oligos for Illumina® Dual Index Primers Set 1 (E7600S). We followed the manufacturer’s protocol for use with inputs ≥100 ng DNA and we used 150 ng of DNA for each library preparation. We aimed for the fragment size range at 200–450 bp; to achieve this we had to reduce the recommended incubation of 15 minutes at 37°C to 11 minutes. We used Beckman Coulter Life Sciences AMPure XP SPRI Reagent (A63881) for the steps requiring magnetic beads and the magnet was ThermoFisher Scientific Invitrogen magnetic stand-96 (AM10027). Our aim for the final size of DNA libraries was approximately 320–470 bp. We verified that and the lack of impurities with Agilent 5200 Fragment Analyzer and ProSize data analysis software. We measured DNA library concentrations with the ThermoFisher Scientific Qubit DS DNA Kit (Q32851).

Once the DNA libraries had passed our quality control, we prepared two pre-pools from individual library samples, one containing three samples with average library size ranging from 320 to 396 bp and the other containing nine samples with average library size ranging from 486 to 579 bp. Equimolar amounts of individual DNA samples were used, based on the library average size data. Finally, we sent these pre-pools on dry ice to Novogene Corporation Inc. (United Kingdom), and they performed the Illumina NovaSeq sequencing runs with 150-base-pair paired-end reads.

We trimmed the raw Illumina reads and adapter sequences with *Trimmomatic* v0.39 (Bolger, Lohse, and Usadel 2014) and then mapped them to a *F. aquilonia* × *F. polyctena* hybrid reference genome (Nouhaud, Beresford, and Kulmuni 2022) with BWA-MEM algorithm in *BWA* v0.7.17 (Li 2013). We removed duplicates with *Picard tools* v2.21.4 (http://broadinstitute.github.io/picard) and clipped overlaps with *BamUtil* v1.0.15 (https://github.com/statgen/bamUtil) These steps had been performed separately also for the previously published data using the same pipeline (Portinha et al. 2022; Satokangas et al. 2023), except for overlap clipping that had not been performed for samples from Portinha *et al*. due to low fraction of read overlap (4% in Portinha *et al*. vs. 13-15% in the other samples). For the previously published data used in this study, average read depth was 16.0× (Satokangas et al. 2023). Our average read depth for the newly sequenced individuals was 10.5× (for individuals that passed all filters and were analysed in this study; see S2 Table), computed with *mosdepth* v0.3.3 (Pedersen and Quinlan 2018).

We performed the following steps for all samples simultaneously. Variants were called for the nuclear genome excluding short genomic regions with no known location (“Scaffold 00” in the genome assembly), with *FreeBayes* v1.3.6, using -k option to disable population priors (Garrison and Marth 2012). We normalised the vcf file with *VT* v0.57721, (Tan, Abecasis, and Kang 2015). We then filtered out sites at two base pair distance from indels, as well as sites that were supported by only forward or reverse reads, with *BCFtools* (Danecek et al. 2021). We decomposed multinucleotide variants using vcfallelicprimitives from *vcflib* v1.0.0_rc3 (Garrison et al. 2022). We filtered the resulting vcf file with *BCFtools* to keep only biallelic single nucleotide polymorphisms (SNPs) with quality equal or higher than 30. We then identified and set missing all sites that had over two times the mean sequencing depth of each individual in question. The excess depth may be an indication of collapsed gene duplications in the reference genome. We further removed sites with heterozygote excess when all samples were pooled, as these sites may represent genotyping errors e.g. due to misaligned reads (*p* < .01, *VCFtools* v0.1.16; (Danecek et al. 2011)). We set individual genotypes that had < 8 coverage as missing.

We excluded seven individuals with more than 50% missing data and three additional *F. rufa* group individuals that were sequenced for collaborative purposes and excluded then sites that had more than 10% missing data over all remaining individuals. We also removed Scaffold 03, the so-called social chromosome with large inversions (Brelsford et al. 2020) as it would bias further computations, and excluded singletons (minor allele count < 2) with *VCFtools*. Extra filtering steps required for some of the analyses that we performed are mentioned with each analysis’ description.

As a result, our dataset (the “main vcf” from here onwards) for this study comprised 93 *F. rufa* group individuals with 1.890.044 biallelic SNPs.

### 2.3. Genetic structure

To confirm the genetic clustering and species identity of the new morphologically identified samples, we recreated the phylogenetic network from Satokangas et al. (2023). Especially, our new *F. rufa* samples helped in confirming whether the closely related and interbreeding sister species *F. rufa* and *F. polyctena* cluster by species regardless of their geographic origin.

Using the main vcf (93 individuals) thinned such that there was at least 20 kb between each variant, resulting in altogether 9.816 biallelic SNPs, we computed pairwise genetic distances of all pairs of samples with distMat.py (github.com/simonhmartin/genomics_general), and generated a phylogenetic network from the resulting distance matrix using *SplitsTree* v4.17.1 (Huson and Bryant 2006) and the NeighbourNet approach (Bryant and Moulton 2004) with default parameters.

### 2.4. Outgroup

To construct and root a species tree and infer introgression in the *F. rufa* group (see below), we used *F. exsecta* as an outgroup. We utilised bam files mapped with the same reference genome as in this study (mapping quality ≥20, sequencing depth 14.5×), generated by Nouhaud et al. (2022). The sequencing data originates from Dhaygude et al. (2019) who had whole-genome sequenced a pool of 50 haploid male ants with 100- base-pair paired-end reads generated on Illumina (ENA accession number SAMN07344806).

We extracted the *F. exsecta* genotypes by calling variants at the biallelic SNP loci with *BCFtools* mpileup (to generate the genotype likelihoods, disabling the computation of base alignment quality and filtering out bases with quality less than 20) and *BCFtools* call (to perform the actual calling), setting ploidy to 1, and removing duplicate sites retaining one allele per locus. This pipeline was adapted from Nouhaud et al. (2022). See each analysis for when the outgroup SNPs were called and merged with the main vcf. The outgroup was merged in all cases after minor allele filtering, as the outgroup is represented only as one haploid genotype per each locus.

### 2.5. Phylogenetics

To investigate the nuclear species tree of the *F. rufa* group, we constructed a neighbour-joining (NJ) tree. The NJ-tree is suitable for very large datasets, and for both within- and between-species trees as it does not force the data with any evolutionary assumptions.

We thinned the main vcf file such that we had a minimum 1 kb distance between each variant with *VCFtools*. We did thinning for both computational reasons but also because in this way regions of the genome which have a distinct history, like those under strong selection, impact the analysis less. We then called the outgroup *F. exsecta* variants and merged the outgroup vcf to the thinned main vcf with *BCFtools* merge. We selected individuals that clustered genomically together by species (S1 Fig, and Satokangas et al. (2023): Fig 1) as including clearly admixed individuals would be unhelpful in constructing the species tree. We further excluded two individuals with more than 30% missing data to maximise the number of SNPs available for the tree, and as the last step excluded all SNPs with more than 1.0% of missing data over all individuals, retaining altogether 73 individuals and the outgroup (See S1 Table), and 165.332 SNPs.

**Fig 1.**
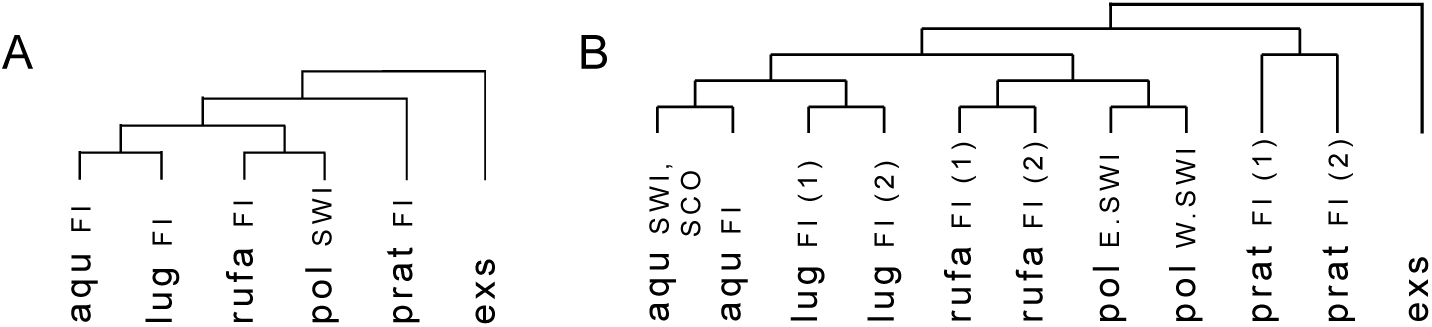
An illustration of the trees used for studying introgression with the *f*-branch statistic. **A** A simple tree with one tree tip per species (5 individuals per species and the outgroup). **B** A tree where each species is split into two groups based on clustering in the species tree (2-3 individuals per species and the outgroup). Species are indicated as aqu = *F. aquiloni*a, lug = *F. lugubris*, rufa = *F. rufa*, pol = *F. polyctena*, prat = *F. pratensis*, exs = *F. exsecta* (used as the outgroup). Origin of sampling is indicated as FI = Finland, SWI = Switzerland, SCO = Scotland, E.SWI = Eastern Switzerland, W.SWI = Western Switzerland.

We then converted the vcf into phylip format with vcf2phylip script (v2.8, (Ortiz, n.d.)) with default parameters and *F. exsecta* defined as the outgroup. We constructed a distance matrix from the phylip file using the *phangorn* package (Schliep 2010) in R, with JC69 (Jukes and Cantor, 1969) substitution model that assumes equal base frequencies and equal mutation rates. We then performed 100 bootstraps with *phangorn*, and visualised the tree using *Figtree* v.1.4.4 (http://tree.bio.ed.ac.uk/software/figtree/) rooting it with *F. exsecta*.

### 2.6. Tests of divergence and diversity

To investigate the genome-wide patterns of speciation, we computed differentiation *F*_ST_ , divergence d_xy_, and nucleotide diversity π in 100 kb genomic windows.

#### 2.6.1. All-sites vcf

For this purpose, we first generated an all-sites vcf file for each scaffold separately with *BCFtools* v1.16 mpileup and *BCFtools* call commands with multiallelic call mode. To calculate a maximum allowed per-site coverage, we utilised the individual mean depths that were computed in the main vcf pipeline for the same purpose. For all-sites vcf, we set the maximum depth per site (over all individuals) as two times the mean depth of per-individual averages. The maximum depth filtering was done to exclude collapsed paralogous genomic regions. We set a minimum depth per site over all individuals as two times the number of individuals (that is, on average, two reads per individual). Sites not meeting these thresholds were set as missing. We then removed the same individuals that we removed from the main vcf, that had shown over 50% missing data, and excluded all sites that had more than 50% missing data. Next, from each scaffold-specific vcf file, we extracted one vcf file with only invariant sites, and another one with only variant sites, with *VCFtools* v0.1.17. In each variant site vcf we kept only sites with minimum quality of 30 and excluded sites that showed excess heterozygosity (vcftools --hwe 0.001). Singletons were filtered out, as rare uninformative markers potentially bias (lower) *F*_ST_ values (Roesti, Salzburger, and Berner 2012). Finally, we combined all scaffolds and the invariant and variant site vcf files with *BCFtools* concat to get one all-sites vcf file and removed duplicate sites. This resulted in an all-sites vcf file with 194.332.157 sites and 93 individuals.

#### 2.6.2. F_ST,_ d_xy,_ and π computation in genomic windows

We computed the *F*_ST_, d_xy_, and π statistics for 100 kb non-overlapping windows, using the *PIXY* package (Korunes and Samuk 2021). *PIXY* was used as it accounts for missing data as this would otherwise lead to biassed estimates. Window size of 100 kb was chosen as it is large enough for liable *F*_ST_ estimates with low sample size (Nadeau et al. ^2^012). Each window contained loci from only one scaffold (no windows crossing scaffold boundaries). To reduce stochasticity, we excluded windows where data for more than half of the sites (50 kb) was missing, as well as windows with less than 100 SNPs. For species pairs’ comparisons, mean π of both species in question was calculated for each genomic window. *F*_ST_ values below zero were converted to zero following a common practice. For *F*_ST_, d_xy_, and π computation we used five individuals per species with least missing data, each from a different population within Finland (as in *f*-branch statistic below). For *F. polyctena* our sampling did not allow this, and we chose five that maximised the number of individuals coming from different populations in Switzerland (see S1 Table).

### 2.7. Tests of introgression

#### 2.7.1. Genome-wide introgression

To test for signatures of introgression between the *F. rufa* group species, we computed the *f*-branch (*f_b_*) statistic with *Dsuite* package (Malinsky, Matschiner, and Svardal 2021). *f_b_* is based on estimating the proportion of genome-wide introgression (*f*) – the relative amount of shared variation between two non-sister taxa P2 and P3, compared to P1 and P3, given altogether four taxa (((P1,P2),P3),Outgroup). *f_b_* helps with interpretation of correlated introgression proportions between related taxa by assigning gene flow not only to tips of a phylogeny but also to the internal branches.

We added the outgroup vcf to our main vcf with *BCFtools* merge. We used the NJ-tree and genetic structure results from this study to provide a tree hypothesis and to group the individuals for the analysis. First, we selected five individuals per species with the least missing data and *F. exsecta* specified as the outgroup (S1 Table) to compute *f_b_* between the species (Fig 1A). This simplest tree, however, did not allow us to evaluate introgression between sister species *F. rufa* and *F. polyctena*, or *F. aquilonia* and *F. lugubris*. Hence, we computed next the *f_b_* statistic for a more detailed tree (Fig 1B), where all species were separated in two groups based on the deepest split in the species tree (Fig 2).

**Fig 2.**
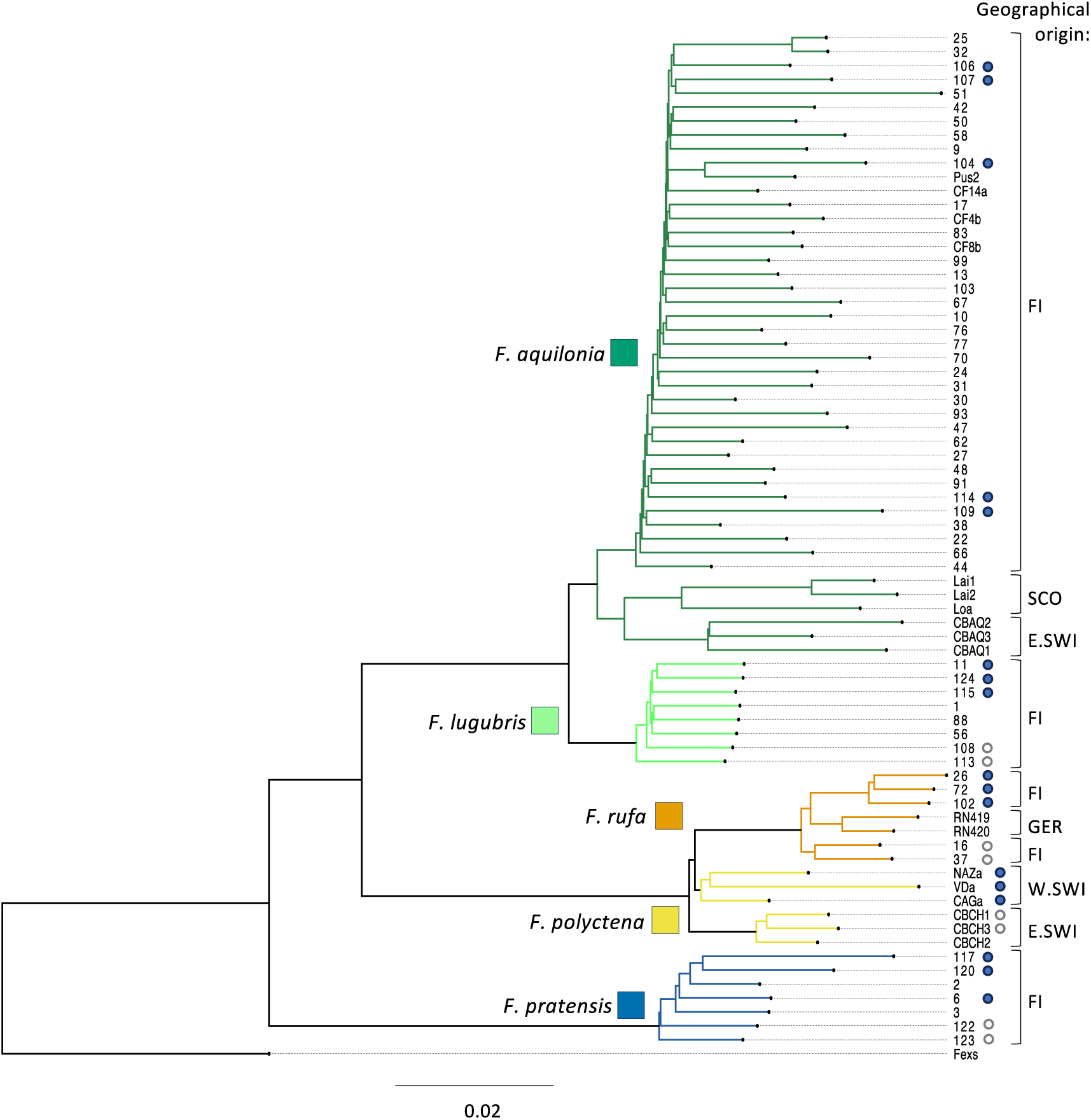
Species tree. Neighbour-joining tree based on whole-genome data (69 individuals and the outgroup, 534.202 SNPs) shows that despite high amounts of unsorted genetic variation, all species but *F. polyctena* form monophyletic groups. The tree is rooted with *F. exsecta* (“Fexs”) and bootstrap support is 100 for all nodes (as percentages of 100 replications). Geographic origin of each individual is shown at the end of the branches. The branch length scale bar indicates the fraction of changed sites. Individuals with blue and white circles were used for all windowed analyses and *f*-branch. The colour difference of the circles shows how the individuals were split in two groups for the second *f*-branch run (Fig 1B). For this second run, part of the *F. aquilonia* individuals were replaced by Swiss and Scottish samples (S1 Table); not shown here.

The motivation for this is that, when introgression is recent or variable in space, individuals of the same species may differ in their proportion of introgression (Hamlin, Hibbins, and Moyle 2020). Such differential introgression can drive phylogenetic clustering, so our decision to use the deepest split within each species to define two groups maximises our potential to detect distinct introgression proportions between the groups. Detection of such differences in introgression between a species and populations of its sister species would in itself be evidence of introgression between these two sister species.

Therefore, the second analysis was run with two to three individuals with the least missing data per group, determined by the availability of individuals per species. If the deepest split separated only one individual from the rest, a second individual was added. For *F. aquilonia*, individuals from Switzerland and Scotland were added for this detailed analysis. For *F. aquilonia* and *F. polyctena*, the within-species split reflected differences in geographical origin.

#### 2.7.2. Introgression in genomic windows

We further investigated the genomic landscape of introgression, especially asking whether introgression is more prevalent in the regions of low diversity and recombination, consistent with signatures of adaptive introgression. For this, we computed the *f_d_* statistic (Martin, Davey, and Jiggins 2015) in 100 kb non-overlapping sliding windows with ABBABABAwindows.py script from https://github.com/simonhmartin/genomics_genera<u>^l^</u>. The *f_d_* is derived from the standard estimator of the proportion of introgression (*f*), applied in *f*-branch. For instance, *f_d_* of 0.1 for a given window indicates that 10% of haplotypes in the recipient population in this genomic window are inferred to be introgressed.

While the standard *f* estimator is sensitive to stochastic errors when the number of studied variants is small (as in windowed analysis), *f_d_* accounts for such stochastic noise by assuming the most conservative direction of introgression at each site (Martin, Davey, and Jiggins 2015). In other words, to compute the standard *f* statistic, the observed excess of shared derived alleles is divided by the expected excess under the maximum possible amount of introgression (complete ‘swamping’). The maximum possible introgression is conventionally estimated by treating P3 (the source population for observed introgression) as both the source and recipient population. When, at a specific site, the assumed recipient population (P2) has a higher frequency of the derived allele than the assumed source population (P3), this may lead to overestimation of windowed *f*-values (*f* may exceed one at a specific site). To correct for this, when calculating the maximum possible introgression, *f_d_* chooses for each site the source population based on which population (P2 or P3) has larger frequency of the derived allele. This eliminates *f* estimates that exceed 1.

We estimated *f_d_* between species pairs for which we identified genome-wide introgression with the *f*-branch statistic, i.e. from *F. polyctena* into *F. aquilonia*, from *F. aquilonia* into *F. polyctena,* and from *F. polyctena* into *F. rufa* (Table 1). Furthermore, we estimated *f_d_* in three species pairs with no significant detected introgression at the genome-wide scale using *f*-branch to confirm that signal of high introgression is not an artefact caused by other genomic properties (Table 1). For studying introgression between *F. polyctena* and *F. rufa,* three individuals per group were used, while for all other comparisons the number of individuals was five. It needs to be kept in mind that the test is always relative, limited to revealing variation shared by P3 and P2, but not P1. Hence, for instance, when inspecting gene flow between *F. rufa* and *F. polyctena*, we are able to detect only introgression that is shared by some *F. rufa* populations (P2) but not the others (P1).

**Table 1.**
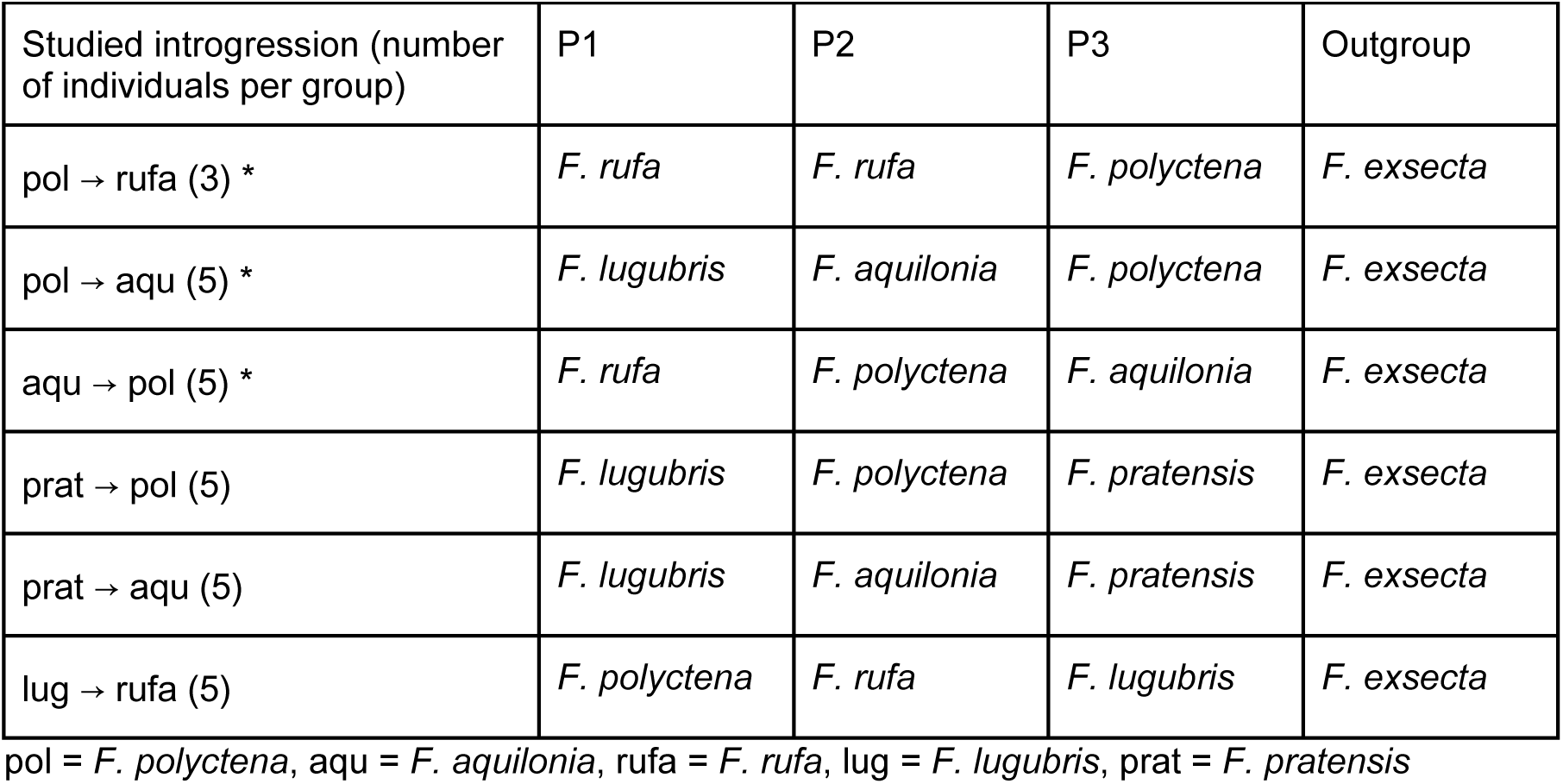
Species combinations used for studying introgression with *fd* statistics in 100 kb windows. * = significant introgression detected with the *f*-branch statistic.

To quantify *f_d_*, windows with nominator values <0 were converted to zero. The nominator quantifies the excess of derived alleles between P3 and P2, and for this statistic, negative values (that would indicate the excess between P3 and P1) are meaningless. Stochasticity of *f_d_* values was minimised by using 100 kb window size and retaining only windows with minimum 100 biallelic SNPs.

### 2.8. Population recombination rate

To investigate the relationship between introgression and recombination (see also section 2.7.2), we used the population recombination rate (rho, ρ) estimates computed in previous work (Nouhaud et al. 2022). The ρ estimates had been computed with *iSMC* software (Barroso, Puzović, and Dutheil 2019) using six *F. polyctena* and six *F. aquilonia* individuals from Switzerland and Scotland (these individuals are also included in this study; see S1 Table), for 20 kb non-overlapping windows. These windows started from varying positions in different scaffolds, which is why we averaged the values over 100 kb non-overlapping windows and 20 kb recombination windows spanning two 100 kb windows were counted in both.

When interpreting the correlations between introgression and recombination or diversity, it needs to be kept in mind that the *iSMC* ρ values and diversity (π) are correlated with each other as both are affected by the local effective population size N_e_. This is because ρ is a composite parameter that is scaled by N_e_ (population recombination rate ρ = 4N_e_*r, where r = recombination rate, and N_e_ is multiplied by four because it accounts for diploidy and recombination events in two lineages), and π is an estimator of the population mutation rate θ that is also scaled by N_e_ (θ = 4N_e_*μ, where μ = per-generation mutation rate) (Watterson 1975; Barroso, Puzović, and Dutheil 2019). These estimators are affected only by those recombination events or mutations that have occurred more recently than the most recent common ancestor of the individuals in question. For this reason, low ρ and low π both occur in regions of low N_e_ regardless of the local crossover rate, which is why observations of low ρ and π may not be independent.

### 2.9. Genome-wide correlations between population genetic parameters

After we had summarised the genomic variation in diversity (π), divergence (d_xy_), differentiation (*F*_ST_), population recombination rate (ρ), and introgression (*f_d_*) in 100 kb non-overlapping sliding windows, we used Spearman’s rank correlation to compute genome-wide correlations between these statistics to investigate the genomic landscape. Spearman’s correlation suits our data as it does not assume linear correlation between variables. We are aware that the 100 kb genomic windows are not independent data points because of genetic linkage of loci between the windows, and that this pseudoreplication may affect the correlation test p-values.

However, the correlation tests provide a valuable way to summarise genomic variation and compare empirical data with theoretical expectations.

To confirm that our correlation results were not affected by higher diversity caused by genomic regions that have been collapsed in the genome assembly and have remained in the data despite vcf filtering, we performed an additional filter step, where we excluded high-coverage outlier windows from our correlation analyses. For this, we computed per-species mean coverage in 100 kb windows from bam files using *mosdepth* v0.3.3 (Pedersen and Quinlan 2018), using the individuals for which the correlation analyses were run. We then removed all windows in which the coverage for any species exceeded two times the mean coverage of that species, thus excluding potentially biassed genomic regions.

We computed genome-wide correlations between a) population recombination rate and introgression (ρ and *f_d_*), b) diversity and introgression (π and *f_d_*), c) diversity and divergence (π and d_xy_), d) diversity and differentiation (π and *F*_ST_), and e) divergence and differentiation (d_xy_ and *F*_ST_), with the following expectations:

a. *Positive correlation between recombination and introgression* is suggested to be indicative of dominating deleterious introgression, with a large number of barrier loci dispersed throughout the genome. Recombination is needed to separate beneficial or neutral introgressing material from the prevalent deleterious material leading to relative excess of introgression in high recombining regions (Veller et al. 2023). In low recombining regions the introgressed material with mostly deleterious haplotypes is selected against. In addition, many alleles with small deleterious effects can be jointly selected against more efficiently than in high recombining regions, as their selective coefficients are combined.

*An absence of correlation or negative correlation* is suggested to arise from neutral or beneficial fitness effects of large introgressing haplotypes, as the haplotypes may get selected for as a whole without the need of partitioning by recombination, or selection on the introgressed neutral haplotypes is not affected by recombination. This may be a consequence of that either polygenic barriers (consisting of many loci) have not yet evolved, or that the direct fitness benefits of the introgressing alleles and coadaptation within the introgressing block outweigh or are equal to the negative epistatic interactions of introgression with the recipient genome.

1. b) *Negative correlation between diversity and introgression* is indicative of adaptive introgression. This suggests that selection has favoured introgression and hence reduced diversity either because introgression transfers adaptive traits or because it may reduce genetic load in the recipient species. However, if both source and recipient species have low diversity in a region of high introgression, then the low diversity (and the correlation) may be caused by strong background selection that has persisted since speciation, rather than by an adaptive introgression event.
2. c) *Strong positive correlation between diversity and divergence* is expected in young species as sequence divergence between young species predominantly reflects ancestral diversity (diversity in the ancestral population that predated the speciation event) or the lack thereof. This correlation weakens over time as sequence divergence accumulates through mutation, drift, and selection. In the face of strong gene flow, divergence may only accumulate in genomic regions where divergent selection acts to limit the effective migration rate, which might eventually lead to a negative correlation between diversity and divergence.
3. d) *The correlation between diversity and differentiation* may be stochastic in early divergence due to sampling effect and differences in sorting ancestral variation, reasoned in a verbal model (Burri 2017). As differentiation increases, the effects of positive selection and later on linked selection (hitchhiking as well as background selection) start to be detectable. A negative whole-genome correlation is expected to arise and strengthen through divergence: selection acts independently in both lineages, and where it has the strongest effect, it may lead to differential fixation (low diversity) and hence elevated differentiation.
4. e) *Positive correlation between divergence and differentiation* is expected in divergence with gene flow, if a large number of gene flow barriers are present throughout the genome.

### 2.10. Local genomic relationship of introgression, and diversity or recombination

We then investigated local patterns of introgression and tested whether there are more introgression outliers than expected in low-diversity, or low-recombining regions. High proportion of introgression associated with low diversity and low recombination is a potential indication that this introgression is favoured by selection.

For this, we first divided the 100 kb genomic windows in bins of low and high diversity, or recombination, by the median value. Then, we calculated the number of top 3%, and 1% windows with highest introgression proportions in both bins and performed a Pearson’s Chi-squared test with Yates’ continuity correction (to correct for potential Type I error when dealing with small expected cell counts) between the top introgression regions and either diversity or recombination.

## 3. Results

### 3.1. Species tree and population structure of the Formica rufa group

To understand the evolutionary relationships of our study species, we constructed a whole-genome neighbour-joining tree using 69 individuals from the five *F. rufa* group species (minimum six individuals per species) and 0.5 million variable sites across the genome, and *F. exsecta* as the outgroup (Fig 2).

The resulting species tree revises the earlier phylogenies (Goropashnaya et al. 2012; Goropashnaya, Fedorov, and Pamilo 2004; Romiguier et al. 2018) (although see (Borowiec, Cover, and Rabeling 2021)) and shows that the *F. rufa* group consists of two sister species pairs *F. aquilonia* and *F. lugubris*, and *F. polyctena* and *F. rufa*, and a basal, more distantly related species *F. pratensis*. Long individual branch lengths relative to internal branches separating the different species indicate high amounts of unsorted genetic variation. Individuals of each species form monophyletic groups with 100% bootstrap support, except for *F. polyctena* that shows nested clustering, which likely suggests geographical differences in the *F. polyctena* introgression history.

We also performed a population structure analysis by constructing a neighbour-joining network for all 93 individuals and ca. 10.000 SNPs, following the pipeline in Satokangas et al. (2023) (S1 Fig). This network demonstrates the relation of newly sequenced samples and the previously published ones, and specifically shows that *F. rufa* and *F. polyctena* cluster by species regardless of their geographic origin, which was an open question based on earlier work.

### 3.2. Genome-wide divergence, differentiation, and diversity

To inspect patterns of genome-wide variation across the group, we studied the levels of divergence (d_xy_), differentiation (*F*_ST_) and diversity (π) between and within the species, computed for altogether 25 individuals (five individuals per species), in 100 kb non-overlapping genomic windows (Fig 3).

**Fig 3.**
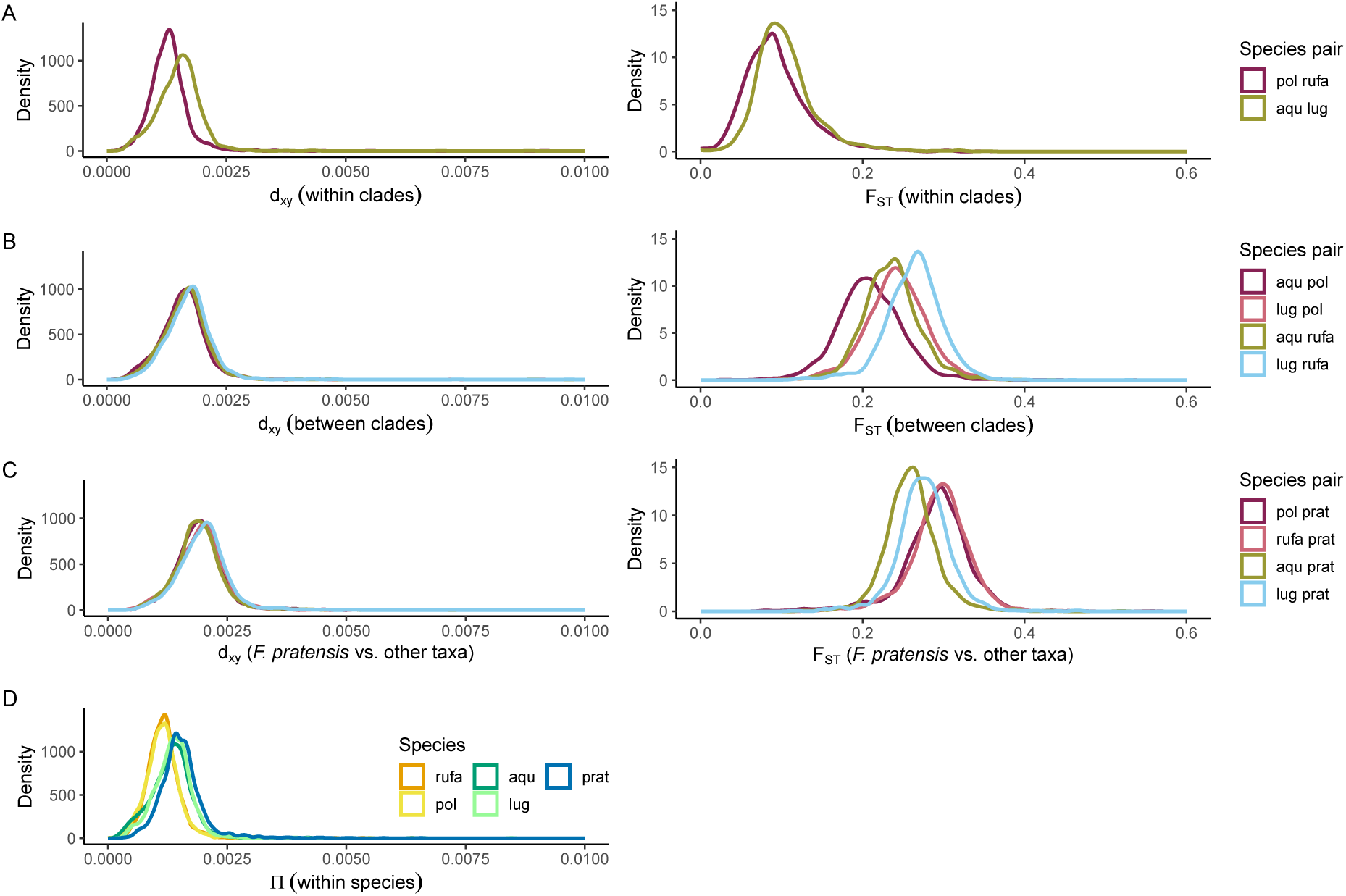
Patterns of genome-wide variation. Levels of divergence (dxy), differentiation (*F*ST), and diversity (π) within the *F. rufa* group computed in 100 kb genomic windows demonstrate early stage of divergence. dxy and *F*ST are compared between **A** sister species, **B** clades (non-sister species), and **C** the basal *F. pratensis* and all other species. **D** presents within-species diversity. Five individuals per species are used, shown in NJ-tree (Fig 2) with circle symbols.

π, the average number of nucleotide differences per site within each species, indicates low levels of diversity across the whole group, with slightly lower values for the southerly distributed species *F. rufa* and *F. polyctena*, in comparison to the others (Fig 3D). d_xy_, the average number of nucleotide differences per site between species, suggests that the group has started to diverge yet the divergence is still low (Fig 3A-C, left panel). This divergence resembles the amount of within-species diversity, particularly in the sister species pairs. *F*_ST_, which reflects the allele frequency differences between the taxa, increases for most part with phylogenetic distance, yet there are differences between species pairs with equally distant phylogenetic histories (Fig 3A-C, right panel).

### 3.3. Correlations between population genetic parameters and inference of genome-wide barriers to gene flow

To help understand the stage of divergence and evolution of gene flow barriers in the *F. rufa* group, we investigated correlations between summary statistics of genetic diversity, divergence, and differentiation using the data computed in 100 kb non- overlapping genomic windows for the 25 chosen individuals (see Table S1) and performing statistical testing with Spearman’s correlation test.

In this group, divergence is highly correlated with within-species diversity (Spearman’s rho = 0.97–0.99, p < 0.001 in all comparisons), as expected for young species in which divergence still reflects ancestral diversity. In all comparisons, divergence is, however, always higher than diversity and this difference increases with phylogenetic distance (Fig 4A, S2 Fig). Differentiation, on the other hand, is only to a very small extent explained by diversity (Spearman’s rho = -0.08 – -0.32, p < 0.001 in all comparisons) and the strength of this correlation is not related to the phylogenetic distance (Fig 4B, S3 Fig). Furthermore, when inspecting the species pairs in which we detect significant genome-wide introgression with the *f*-branch statistic (*F. polyctena* and *F. rufa*, and *F. polyctena* and *F. aquilonia*, see 3.4.), we find no genome-wide correlation between divergence and differentiation (Fig 4C, S4 Fig), indicating that barriers to gene flow are not yet strong enough to produce a genome-wide signal of positive correlation. Alternatively, as divergence still reflects ancestral diversity, it may be that genome-wide dispersed gene flow barriers do exist, but there has been insufficient time for divergence to develop a detectable signal.

**Fig 4.**
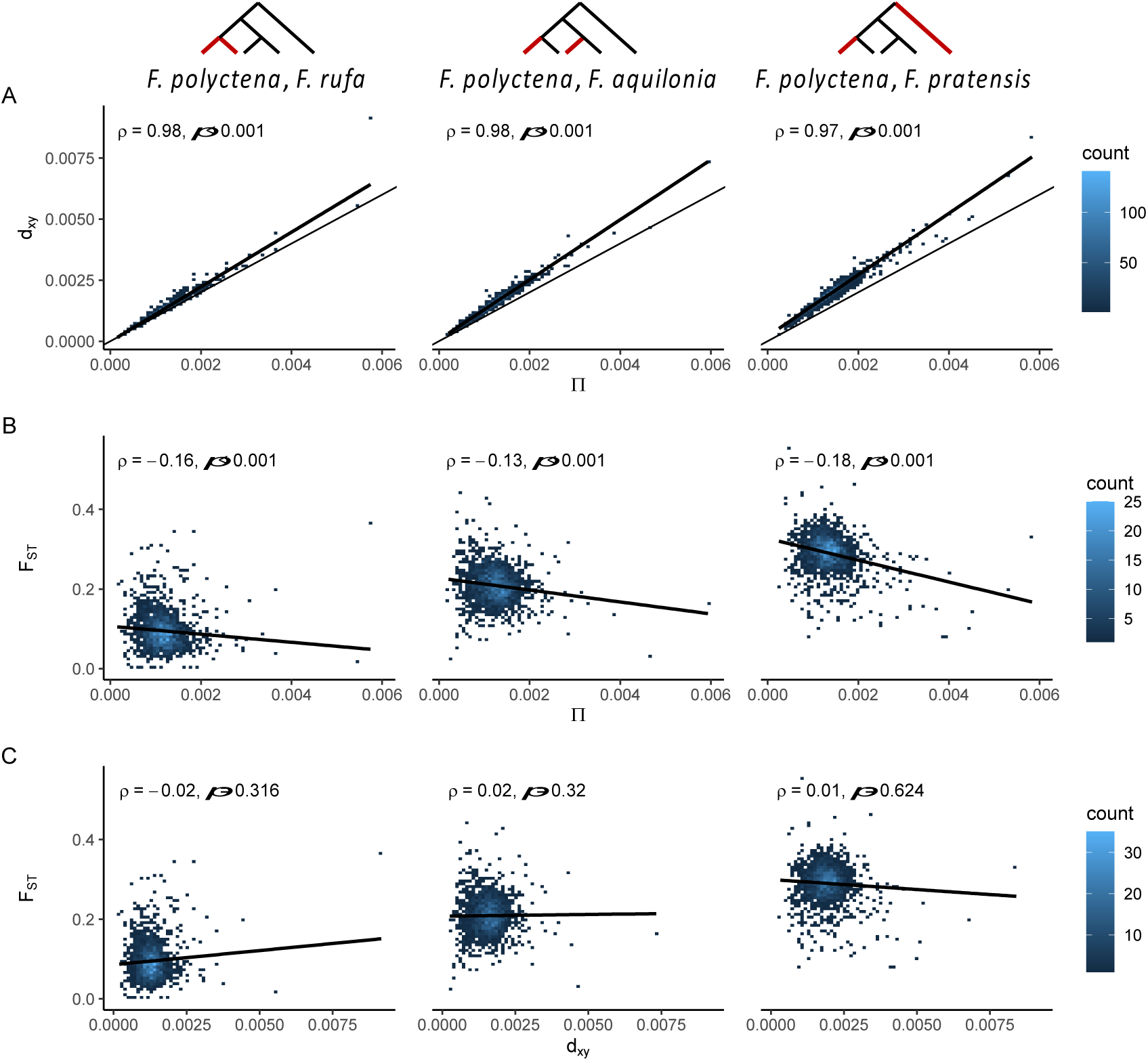
Genome-wide correlations between population genetic parameters along the early stages of speciation continuum. Correlations between population genetic parameters are shown for three species pairs with increasing phylogenetic distance (for the rest of the comparisons, see S2-S4 Fig). The values are computed in 100 kb genomic windows. **A** Divergence (dxy) is highly correlated with diversity within species (π, averaged for the species compared) in all species pairs. dxy is higher than average π in all species pairs and this difference strengthens with phylogenetic distance. Diagonal line with slope 1 is drawn to help see how dxy and π relate. **B** Small and similar proportion of differentiation (*F*ST) is explained by diversity (π), and this pattern does not change despite increasing phylogenetic distance between species pairs. **C** As differentiation (*F*ST) between species pairs increases so does divergence (dxy), however, there is no indication of positive genome-wide correlation in the introgressing species pairs (left and middle column). Correlation tests are performed with Spearman’s correlation. A linear regression line with its 95% confidence interval (shading) is shown.

### 3.4. Introgression at genome-wide and local scale

We first examined signals of genome-wide introgression using the *f*-branch statistic, using 1.9 million genome-wide SNPs and the chosen 25 individuals (five per species), supplemented with six *F. aquilonia* individuals from different geographic locations to allow a more detailed analysis.

According to the most parsimonious interpretation, we found evidence for introgression between two species pairs: sister species *F. polyctena* and *F. rufa*, and non-sister species *F. aquilonia* and *F. polyctena* (Fig 5 A-C). In *F. aquilonia* and *F. polyctena*, the introgression was inferred to be bidirectional (Fig 5A and Fig 5B: i),ii), and iii)). For *F. polyctena* and *F. rufa,* the detected excess of introgression between *F. polyctena* and part of the *F. rufa* populations suggests introgression at least from *F. polyctena* into *F. rufa* (Fig 5B: iv)), while bidirectional introgression cannot be ruled out.

**Fig 5.**
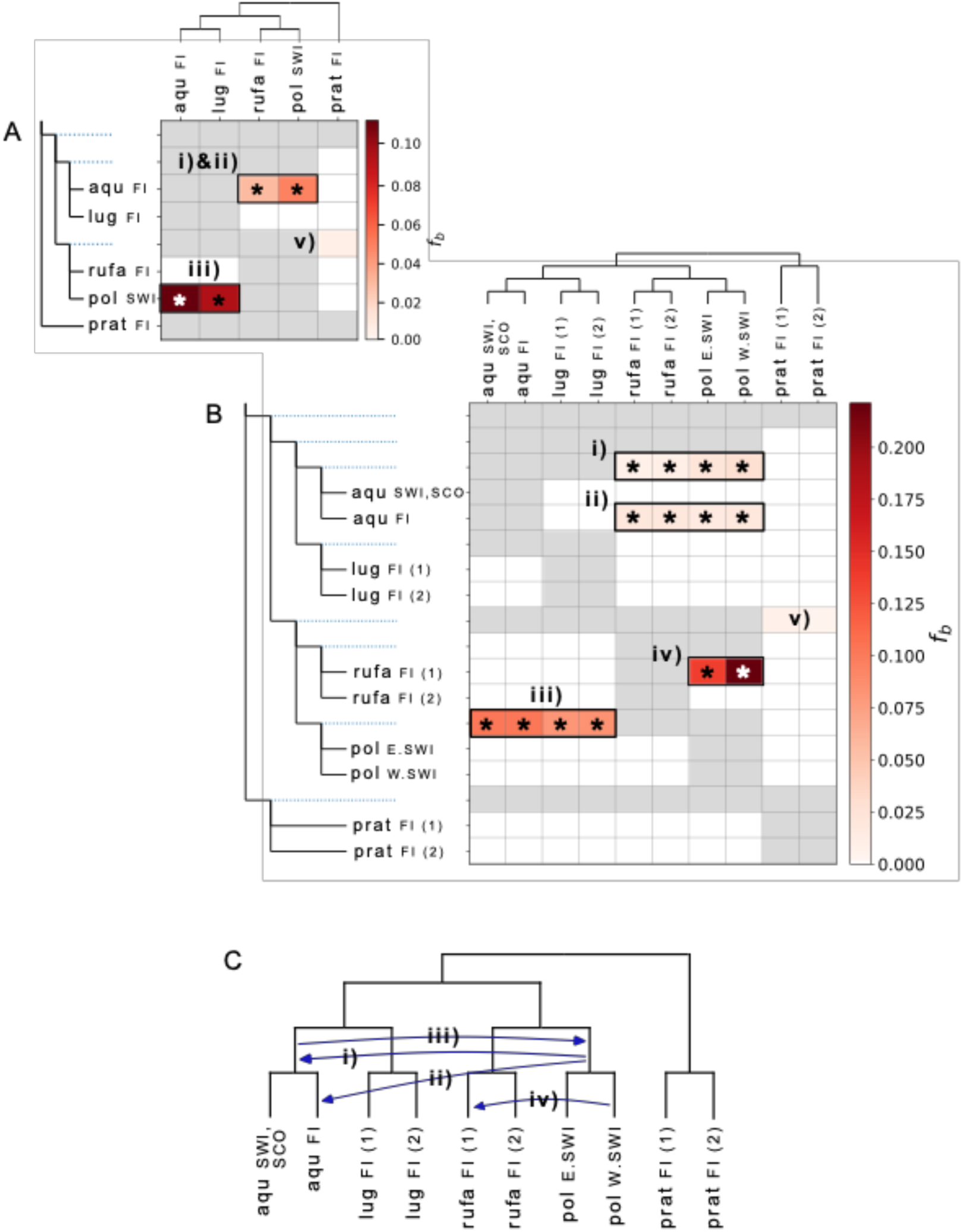
Introgression between species. *f*-branch statistic demonstrates introgression in at least two species pairs. **A** The simplest scenario suggests bidirectional introgression between *F. polyctena* and *F. aquilonia* (i)&ii), and iii)). Inference of ancestral or within-clade introgression is not possible in this tree with one tip per species. **B** *f*-branch statistic computed for a tree, where each species is split according to clustering in the species tree reveals additional introgression events and provides more detailed information. Following the most parsimonious interpretation i) introgression from *F. polyctena* has occurred into the ancestor of Swiss and Finnish *F. aquilonia* populations, and ii) it has continued into the Finnish *F. aquilonia* after the geographical split of *F. aquilonia* populations. Additional introgression has occurred from *F. polyctena* into *F. rufa* (iv)). v) Potential signal of introgression between *F. pratensis* and the *F. polyctena/F.rufa* clade is slightly below significance threshold. Grey data points indicate tests that were not possible given the phylogeny. The horizontal patterns may arise from correlated ancestries between sister taxa of the introgression source, which provides the most parsimonious explanation for the introgression events. Introgression between *F. polyctena* and the ancestor of the two *F. rufa* groups cannot be tested but is likely given the extent of their hybridisation Europe-wide. Asterisks (*) indicate significant test results (z-score >3). **C** A graphical summary of the inferred most parsimonious introgression events.

More specifically, the results suggest that introgression from *F. polyctena* into *F. aquilonia* has occurred into the ancestor of Swiss and Finnish *F. aquilonia* populations and continued into the Finnish but not Swiss *F. aquilonia* after their geographical split. The z-score for introgression between *F. pratensis* and the clade of *F. polyctena* and *F. rufa* (2.8 and 2.7 in Fig 5A and 5B, respectively) remained slightly below the significance threshold (z-score 3).

We then investigated genome-wide correlations between the estimated rate of introgression (*f*_d_), and diversity or recombination in 100 kb windows, in those species pairs for which significant genome-wide introgression was inferred with *f*-branch. This was performed to see whether genome-wide introgression patterns indicate prevalence of deleterious, neutral, or adaptive introgression.

At the whole-genome level, we detected non-existent or a very weak positive correlation between the estimated extent of introgression and both diversity (Fig 6A) and recombination (Fig 6B). This suggests that contrary to the conventional view (Veller et al. 2023) but according to new expectations (Dagilis and Matute 2023), introgression may not be predominantly deleterious in young species.

**Fig 6.**
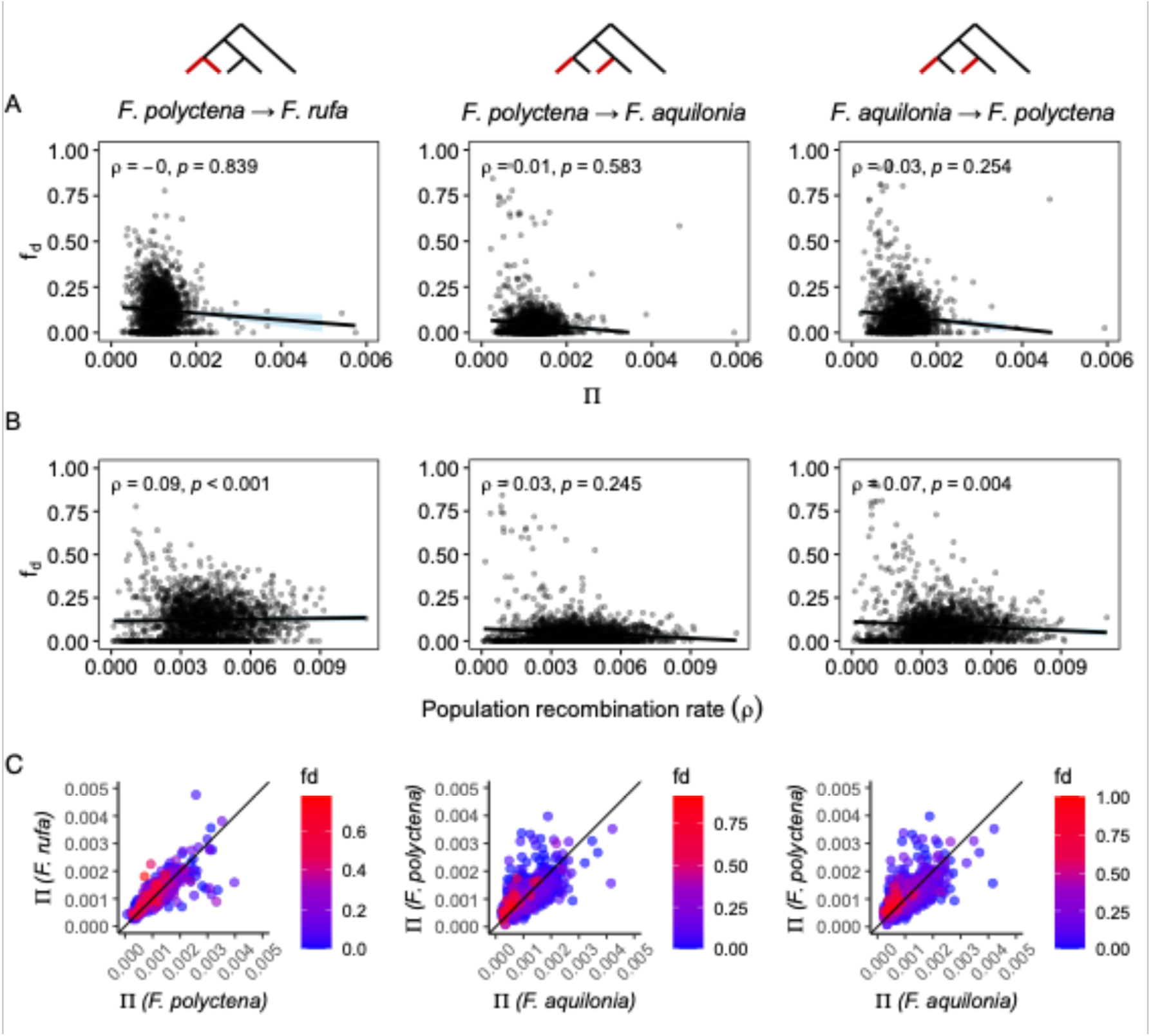
Genome-wide correlations between introgression (*f*d), diversity (π), and recombination (ρ). In species pairs with detected significant introgression, genome-wide correlation between introgression and diversity or recombination is non-existent or very weakly positive. Computed in 100 kb windows. **A** Correlation between diversity (average of the two species) and introgression, **B** Correlation between population recombination rate (average of *F. aquilonia* and *F. polyctena*) and introgression, **C** Correlation between diversities within each species, coloured with the proportion of introgression. Diagonal line with slope 1 helps to compare the diversity between gene flow source and recipient species in regions with high proportion of introgression in panel C. Correlation tests are performed with Spearman’s correlation. A linear regression line with its 95% confidence interval (shading) is shown in panels A and B.

Furthermore, we tested statistically, whether high local proportions of introgression are associated with low diversity or low recombination as such association could be a signal of adaptive introgression. The relation between high proportion of introgression and both diversity (averaged for the species in question) and recombination were significant in both *F. aquilonia* - *F. polyctena* species pair, and *F. polyctena* - *F. rufa* species pair, when both 1% and 3% top introgression outlier windows were examined (Table 2). High introgression was always associated with lower diversity and lower population recombination rate than expected by chance. In addition, to examine the underlying reason behind the association of low diversity and high introgression, we examined visually species-specific diversity in relation to estimated introgression and inferred that the highest estimated proportion of introgression is often located in regions with relatively low diversity in both the introgression source and recipient species, although variation in this does exist (Fig 6C).

**Table 2.** Association of high introgression, and diversity or recombination. Locally in the genome, high proportions of introgression are associated with low diversity and low recombination. The data is computed in genomic windows of 100 kb and tested with chi-squared test, N(aqu pol)=1930, N(pol rufa)=1909. pol = *F. polyctena*, aqu = *F. aquilonia*, rufa = *F. rufa.* * = significant at 0.05 level, ** = significant at 0.005 level

| Introgressing species and the direction of gene flow | % of top outlier introgression windows ($f_d$ threshold) | Count of the outlier windows in each bin | | Diversity ($\pi$) | | | Population recombination rate ($\rho$) | | |
| --- | --- | --- | --- | --- | --- | --- | --- | --- | --- |
| | | $\pi$: low/high | $\rho$: low/high | $\chi^2$ | df | p-value | $\chi^2$ | df | p-value |
| pol → aqu | 1% (0.438) | 16/4 | 19/1 | 6.1249 | 1 | **0.013 \*** | 14.619 | 1 | **< 0.001 \*\*** |
|  | 3% (0.165) | 49/9 | 47/11 | 27.079 | 1 | **< 0.001 \*\*** | 21.813 | 1 | **< 0.001 \*\*** |
| aqu → pol | 1% (0.535) | 19/1 | 19/1 | 14.619 | 1 | **< 0.001 \*\*** | 14.619 | 1 | **< 0.001 \*\*** |
|  | 3% (0.308) | 49/9 | 48/10 | 27.079 | 1 | **< 0.001 \*\*** | 24.375 | 1 | **< 0.001 \*\*** |
| pol → rufa | 1% (0.466) | 15/5 | 19/1 | 4.0929 | 1 | **0.043 \*** | 14.603 | 1 | **< 0.001 \*\*** |
|  | 3% (0.358) | 41/18 | 38/21 | 8.4652 | 1 | **0.004 \*\*** | 4.4774 | 1 | **0.034 \*** |

Finally, we inspected whether the association of high introgression (*f*_d_) and low diversity or recombination is a true signal or if it may be explained by increased noise. Population recombination rates and diversity estimates are lowered by low effective population size (N_e_). In these low N_e_ regions strong drift may create high *f*_d_ values that are not real. Hence, a false association between high *f*_d_ and low π or low ρ may occur because of low N_e_. Additionally, in regions with low actual recombination rate, the level of independence among loci is low, which may increase stochasticity in *f*_d_ values (Martin, Davey, and Jiggins 2015).

To test whether the association of high introgression and low diversity can be explained by noise alone, we first inspected the levels of *f*_d_ in three species pairs with no detected significant genome-wide introgression. Equally high levels of inferred introgression in these control pairs in regions of low π or ρ in comparison to species pairs with significant introgression would indicate that *f*_d_ is elevated because of noise. However, the overall mean *f*_d_, and specifically the local signal of high introgression is lower in control species pairs (S5 Fig), signalling that the high *f*_d_ values cannot be explained by noise alone. Second, we inspected the association between low negative *f*_d_ values, and diversity (π) or recombination (ρ). The motivation is that the association of low ρ and both negative and positive outlier *f*_d_ values may be caused by increased noise in *f*_d_, while a similar excess of negative *f*_d_ as positive outliers is not expected in regions of low ρ if the signal is true. Such noise can arise due to increased drift in regions of low local N_e_ and nonindependence of genomic loci among low recombining regions. Our results demonstrate that locally in the genome low ρ is associated with low negative *f*_d_ values, indicating that the association of low ρ and high outlier *f*_d_ values is potentially caused by noise. The association of diversity and outlier *f*_d_ regions cannot, however, be explained by noise alone. The genome-wide correlations between *f*_d_ and diversity or ρ remained the same despite the inclusion of negative *f*_d_ values, overruling the possibility that high *f*_d_ values caused by noise would have affected our whole-genome correlations (S6 Fig, S3 Table).

## 4. Discussion

The role of introgression in adaptive evolution is increasingly acknowledged across taxa with different levels of divergence (Edelman and Mallet 2021). Recent theoretical work revisits the conventional expectation that introgression may only remain in the genome in regions of high recombination, seen as a positive correlation between recombination and introgression. This expectation arises because introgression is assumed to be mostly deleterious and is therefore selected against in low recombining regions, while in high recombining regions, recombination can separate small adaptive or neutral parts from the introgressed material, letting them to be maintained in the population (Veller et al. 2023). The recent work suggests that in young species large introgressing blocks may remain intact because they may instead have positive or neutral fitness effects (Dagilis and Matute 2023). Using whole-genome resequencing data from *Formica rufa* group we find at most a weak positive genome-wide correlation between introgression and recombination despite polygenic barriers described in earlier work (Kulmuni and Pamilo 2014; Kulmuni et al. 2020). This contrasts with findings from more diverged taxa with relatively less introgression in low recombining regions (Martin et al. 2019; Schumer et al. 2018). Hence, our results are in line with the novel theoretical expectations (Dagilis and Matute 2023). Moreover, we identify local genomic signatures of introgression that has likely been favoured by selection. When hybridization and introgression have both benefits and costs, their long-term impacts for species are challenging to predict. Our findings shed light on the role of introgression in a taxon group, where previous research has inferred both polygenic gene flow barriers and deleterious fitness consequences of hybridisation (Kulmuni and Pamilo 2014; Kulmuni et al. 2020), but also potential adaptive benefits of hybridisation (Martin-Roy et al. 2021; Satokangas et al. 2023; Nouhaud et al. 2022).

### 4.1. No strong genome-wide signature of deleterious introgression

Our data suggests there is at most a weak positive genome-wide association between recombination and introgression. This is in line with the recent theoretical prediction that when the overall divergence is low, the fitness benefits of large introgressing blocks may outweigh their fitness costs. These fitness benefits may arise from direct fitness advantage of the introgressed material, or from preserving together the coadapted alleles of the introgressed haplotype. Earlier research demonstrates this, for instance, in *Drosophila*, where relatively more introgression in regions of low recombination was observed in secondary contact (Duranton and Pool 2022; Pool 2015; Dagilis and Matute 2023). Similarly, research on primates suggests a lack of correlation between recombination and introgression in recently diverged species pairs (Jensen et al. 2023).

We inferred introgression between two species pairs; the sister species *F. rufa* and *F. polyctena*, and non-sister species *F. aquilonia* and *F. polyctena*. Hybridisation between both species pairs is prevalent in nature (Seifert 2021; Seifert, Kulmuni, and Pamilo 2010; Satokangas et al. 2023). There are two plausible reasons that can explain the very weak, or lack of positive genome-wide correlation between introgression and recombination found here. First, it could be that the polygenic barriers have not evolved yet. Barriers to gene flow that would be present across the genome could result in selection against large introgressing blocks and hence selection against recombination in low recombining regions. This may be likely regarding the closely related sister species *F. rufa* and *F. polyctena*. However, it is contrary to earlier findings of polygenic barriers in the non-sister species *F. aquilonia* and *F. polyctena* (Kulmuni and Pamilo 2014; Kulmuni et al. 2020), in which recessive incompatible loci scattered across the genome were shown to cause hybrid male mortality (Kulmuni, Seifert, and Pamilo 2010; Kulmuni and Pamilo 2014). In the non-sister species, perhaps a more likely explanation would be that the benefits of introgression outweigh the fitness costs of genetic incompatibilities that are revealed in introgression. Introgression has a beneficial fitness effect if it delivers adaptive alleles (Edelman and Mallet 2021), masks and helps purging deleterious mutations like in brook charr (Leitwein et al. 2019) and humans (Harris and Nielsen 2016), or when coadaptation within introgressing haplotypes is costly to break, as shown by theoretical work (Dagilis and Matute 2023). We consider the alternative that the benefits of introgression outweigh their costs more likely at least in the case of the non-sister species *F. aquilonia* and *F. polyctena*.

Interestingly, natural selection is unstable at some of the barrier regions between *F. aquilonia* and *F. polyctena,* switching from acting against hybridisation to favouring it across a 10-year period and pointing towards environment-dependent incompatibilities (Kulmuni et al. 2020; Martin-Roy et al. 2021). We do not know yet how such instability affects the fitness effects and signatures of introgression. However, this question is recognised in recent research, and for instance in flowering plants, environmental fluctuations are suggested to affect the amount of gene flow and reproductive isolation between recently diverged subspecies (Sianta, Moeller, and Brandvain 2023).

### 4.2. Signatures of positive selection in regions of high introgression and the consequences of multi-species introgression

Recent research has provided evidence that introgression is an important source of genetic variation in adaptive evolution (Edelman and Mallet 2021). We detected low nucleotide diversity in regions of high introgression, suggesting introgression has been favoured locally in the genome in both the sister and non-sister species pairs. Relatively low diversity was found in both the introgression source and recipient species, suggesting that both the transfer of adaptive alleles and purging or masking of genetic load (deleterious recessive alleles) would be biologically reasonable causes. Earlier work in wood ant hybrids supports at least the genetic load scenario. Populations of hybrids between the non-sister species *F. aquilonia* and *F. polyctena* show predictable ancestry sorting, in which genetic load in the species with lower effective population size, *F. polyctena*, seems to play a role (Nouhaud et al. 2022).

Eusocial Hymenoptera typically have low effective population sizes (Weyna and Romiguier 2021), which may lead to accumulation of genetic load (Kim, Huber, and Lohmueller 2018). Our results show low genome-wide nucleotide diversity in all studied species, which may be a manifestation of low effective population size in the *F. rufa* group. Even if both the introgression source and recipient species have low effective population size that causes accumulation of deleterious mutations, these mutations may occur in different sites and introgression may be beneficial, as demonstrated in humans (Harris and Nielsen 2016). The long-term impacts of such a process are, however, difficult to predict. The once beneficial and favoured Neanderthal introgression has turned out later to be costly due to prevalence of genetic load it brought along to modern humans, and has been in the long term selected against (Schumer et al. 2018; Edelman and Mallet 2021).

Introgression may, on the other hand, represent a source of locally adaptive alleles. It has been suggested that the hybrids between the non-sister species *F. aquilonia* and *F*. *polyctena* may benefit from combining the climatic adaptations from the divergently adapted parental species. This may help them cope with a changing climate (Martin-Roy et al. 2021; Satokangas et al. 2023). From the perspective of the hybridising species, the northern cold-adapted *F. aquilonia* is suggested to suffer from warm winters (Sorvari, Haatanen, and Vesterlund 2011) and hence could benefit if introgression from *F. polyctena* mediated tolerance towards warm conditions. Introgression with adaptive potential in the context of climate change is topical and has been detected for instance in red oaks, in which it is associated with drought tolerance and influenced by habitat conditions (Khodwekar and Gailing 2017) Other reasons for the local association of low diversity and high introgression than adaptive introgression need to be considered as well. Signal of introgression may be biassed by stochastic fixation due to both linked variation and low effective population size. Extensive simulations (Martin, Davey, and Jiggins 2015) show that the *f*_d_ statistic used here outperforms other comparable *f*-statistic estimators in reliability when recombination is low. However, we cannot fully rule out that stochastic fixation would explain the signal of adaptive introgression, seen as the high local introgression estimates in regions of low recombination and low diversity. Additionally, low diversity in both the recipient and the source species in a region of high introgression might be caused by long term strong background selection instead of adaptive introgression. These alternative explanations will need to be addressed in future work. Our current approach with 100 kb genomic window size does not reveal fine-scale variation in introgression and diversity along the genome. Hence, to reveal potential drivers of high introgression in regions of low diversity, it will be essential to study these genomic regions in finer detail.

Sometimes introgression may involve more than two taxa. These so-called interbreeding complexes have been reported in a variety of organisms from plants to invertebrates and vertebrates (see, e.g., Grant and Grant (2020) and references therein), but their evolutionary consequences are poorly understood. Our results show that the wood ants form an interbreeding complex, as the species pairs in which we detected introgression share a common partner. The introgressing alleles may potentially pass through all three species. Our data suggests that if introgression is transmitted across all three species, it could flow from northerly distributed, cold-adapted *F. aquilonia* through southerly distributed, warm-adapted *F. polyctena*, to southerly distributed *F. rufa*. The other direction (from *F. rufa* through *F. polyctena* to *F. aquilonia*) could also be possible, given bidirectional gene flow between the non-sister species and earlier inferences of gene flow in the opposite direction in the sister species (Seifert, Kulmuni, and Pamilo 2010). Hence, gene flow between multiple taxa could allow alleles to be transferred between taxa that do not otherwise hybridise. We cannot exclude the possibility of direct introgression also between *F. aquilonia* and *F. rufa*, as hybrid populations between them exist (Satokangas et al. 2023). However, our results indicate that if such introgression occurs, it is not as strong as that between the other species pairs. We do not yet know the consequences of this triad hybridisation and introgression. There are examples of how introgressing species may act as “genetic bridges” that deliver and hence increase genetic variation across taxa, as seen in the Galapagos finches (Grant and Grant 2020), and of how hybridisation among multiple lineages may facilitate extremely fast adaptive radiations, as seen in African cichlids (Meier et al. 2023).

### 4.3. Differentiation and divergence follow patterns expected for young species

Our results are in line with expectations for young species, as we detect low overall divergence, high positive correlation between divergence and diversity, and weak correlation between differentiation and diversity (Burri 2017). Other closely related taxa, like the black-and-white flycatchers, show similar divergence, differentiation, and diversity estimates as well as similarly strong positive correlation between diversity and divergence (Chase, Ellegren, and Mugal 2021). In addition, in the wood ants, the extensive hybridisation detected previously (Satokangas et al. 2023) and the high amount of unsorted genetic variation revealed by our NJ-tree are typical for young taxon groups.

Our data indicated that gene flow barriers do not manifest as a positive whole-genome correlation between divergence (d_xy_) and differentiation (F_ST_) in this group. When speciation occurs with gene flow, genomic regions that restrict the gene flow are expected to show both elevated differentiation and elevated divergence (Cruickshank and Hahn 2014). In the presence of polygenic gene flow barriers, a positive genome-wide correlation between divergence and differentiation would be expected (Shang, Field, et al. 2023). Hence, such correlation could have been likely especially in the non-sister species *F. aquilonia* and *F. polyctena* that have diverged with gene flow (Portinha et al. 2022) and have polygenic gene flow barriers (Kulmuni and Pamilo 2014; Kulmuni et al. 2020). The first reason for the lack of correlation could be that the barriers are not strong enough, which may be unlikely given the polygenic barriers and hybrid mortality (Kulmuni and Pamilo 2014; Kulmuni et al. 2020). A likely alternative could be that strong barriers exist but divergence has not had enough time to develop a detectable signal in the barrier regions. This explanation is supported by the fact that we found genome-wide divergence to still resemble ancestral diversity in the whole *F. rufa* group.

Previous work has presented opposing views about the phylogenetic relationships in the *F. rufa* group. The species tree we constructed in this study consists of two sister species pairs and a more basal *F. pratensis.* It is in line with a recently constructed mitochondrial network (Satokangas et al. 2023) and an earlier nuclear phylogeny that contains four out of our five study species (Borowiec, Cover, and Rabeling 2021). However, it revises other formerly constructed mitochondrial (Goropashnaya et al. 2012; Goropashnaya, Fedorov, and Pamilo 2004) as well as nuclear (Romiguier et al. 2018) phylogenies. These likely suffer from low sample size (from one to maximum a few individuals per species) that may result in incorrect phylogeny if the individual used happens to have hybrid origin; a plausible possibility given the high hybridisation in the *F. rufa* group (Seifert 2021; Satokangas et al. 2023). The monophyly of *F. rufa* despite sampling from different geographic locations confirmed that it forms a distinct gene pool from its sister species *F. polyctena,* which remained open in previous work (Satokangas et al. 2023).

## 5. Conclusions

Using whole-genome sequencing data, we have inspected divergence and introgression in extensively hybridising wood ants. We showed that there is no strong genome-wide signature of deleterious introgression – a finding that concerns both the sister and non-sister species in this young species group. This was revealed by at most a weak correlation between recombination and introgression, which is in line with recent theoretical work. We also detected local signatures of positive selection acting for the introgressed material. Previous work in wood ant hybrids gives reason to suggest that introgression could have potential to help the *F. rufa* group species cope with genetic load or facilitate their survival in changing climate. Further work on verifying and investigating the signature of adaptive introgression in more detail will shed light on the evolutionary impacts of introgression in the wood ants. This work highlights the notion that evolution towards full reproductive isolation may not promote species persistence in the long run. Instead, divergent taxa may gain by not completing speciation (Barton 2020), although the long-term impacts of introgression are complex to predict.

## Acknowledgements

We thank Airi Lamminmäki for helping with DNA extractions and library preparations for the new ant samples. We are grateful to Pierre Nouhaud, Sean Stankowski, Ari Löytynoja, Patrick Heidbreder and the whole SpecIAnt group, and Simon Martin’s lab, for insightful discussions. The expertise and work of Roland Schultz was essential for having the background data for morphological analyses.

## Data availability statement

All FASTQ files will be available on ENA, and VCF files, windowed statistics and bioinformatic scripts will be available on Dryad and Zenodo upon publication in a peer-reviewed journal. The bioinformatic scripts are currently available at https://github.com/isatok/Wood_ant_local_genomic_patterns.

