## Supplementary material for "Introgression and divergence in a young species group": S1 Fig

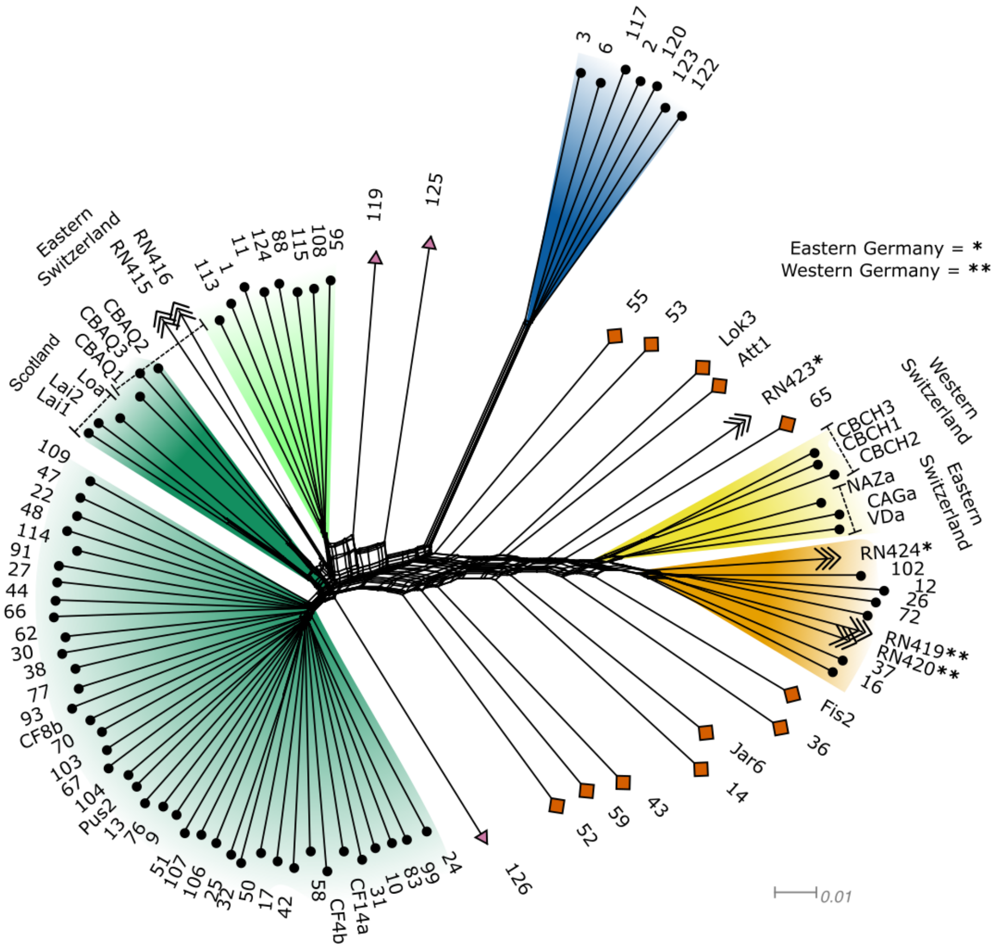


**Fig S1. Neighbour-joining network** performed with 9.816 variants, following pipeline in (Satokangas et al. 2023) to demonstrate how new sequenced samples relate with previously published ones. Arrowheads indicate newly sequenced individuals. The network especially shows how *F. rufa* and *F. polyctena* cluster by species regardless of geographic origin. All individuals for which geographical location is not given originate from Finland. Newly sequenced individuals RN415, RN416, and RN423 are potentially admixed given their placement in this network. Colours indicate species assignments based on combined morphological and genetic data from this study and previous work (Satokangas et al. 2023). Dark green = *F. aquilonia*, light green = *F. lugubris*, dark blue = *F. pratensis*, yellow = *F. polyctena*, orange = *F. rufa*, red diamond = admixed individual between *F. aquilonia* and the *F. polyctena/F. rufa* clade, pink triangle = admixed *F. lugubris* individual.
