## Supplementary material for "Introgression and divergence in a young species group": S2 Fig

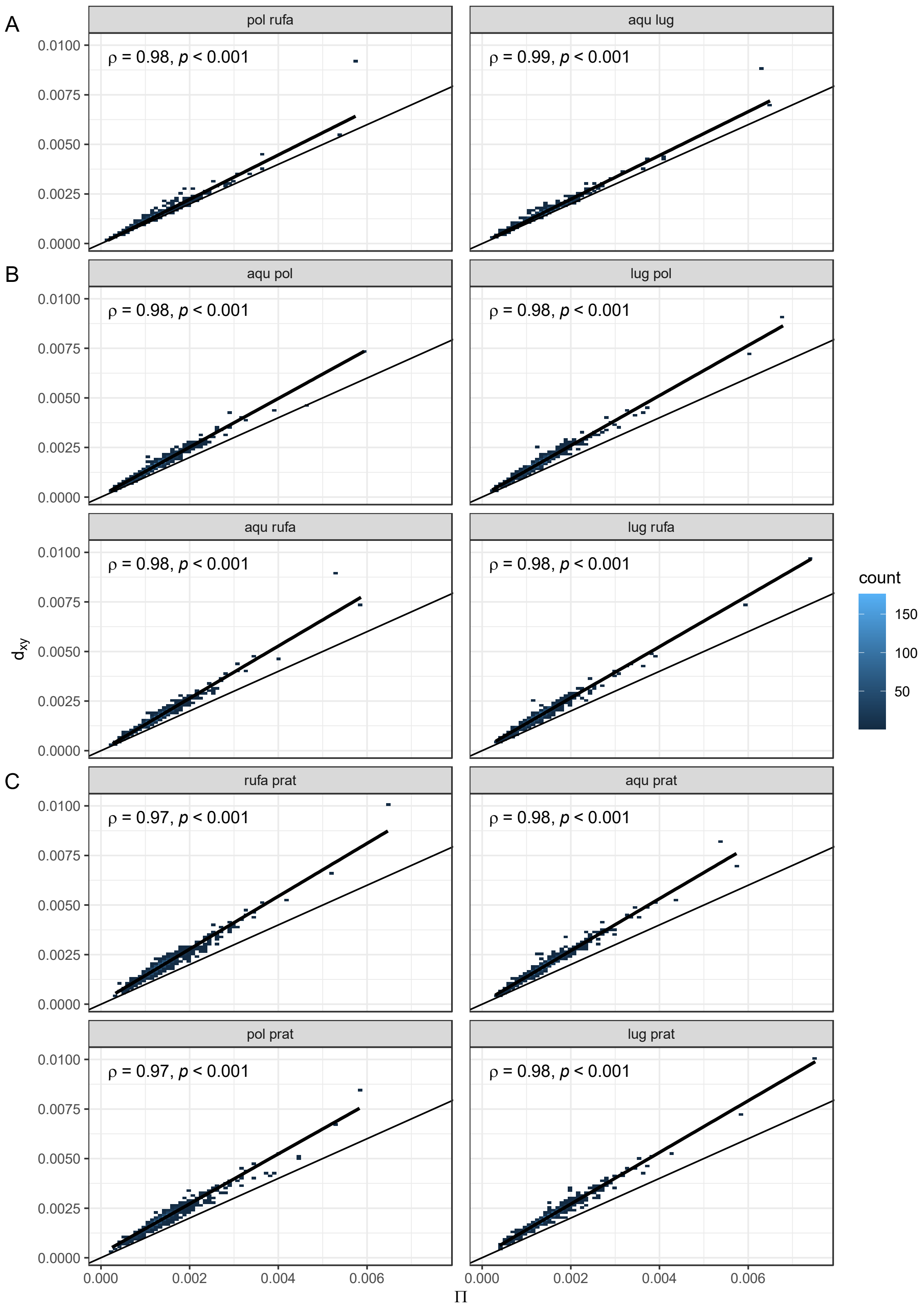


**Fig S2. Correlations between diversity (π) and divergence ($d_{xy}$).** Strong positive correlation between diversity (x-axis) and divergence (y-axis) in all species pairs, computed in 100 kb genomic windows. Diagonal line with slope 1 demonstrates how divergence is higher than diversity, and the difference increases with phylogenetic distance. **A** Sister species pairs, **B** Between-clade species pairs, **C** All species compared to *F. pratensis.* Species are indicated as aqu = *F. aquiloni*a, lug = *F. lugubris*, rufa = *F. rufa*, pol = *F. polyctena*, prat = *F. pratensis.* Correlation tests are performed with Spearman's correlation. A linear regression line with its 95% confidence interval (shading) is shown.
