## Supplementary material for "Introgression and divergence in a young species group": S4 Fig

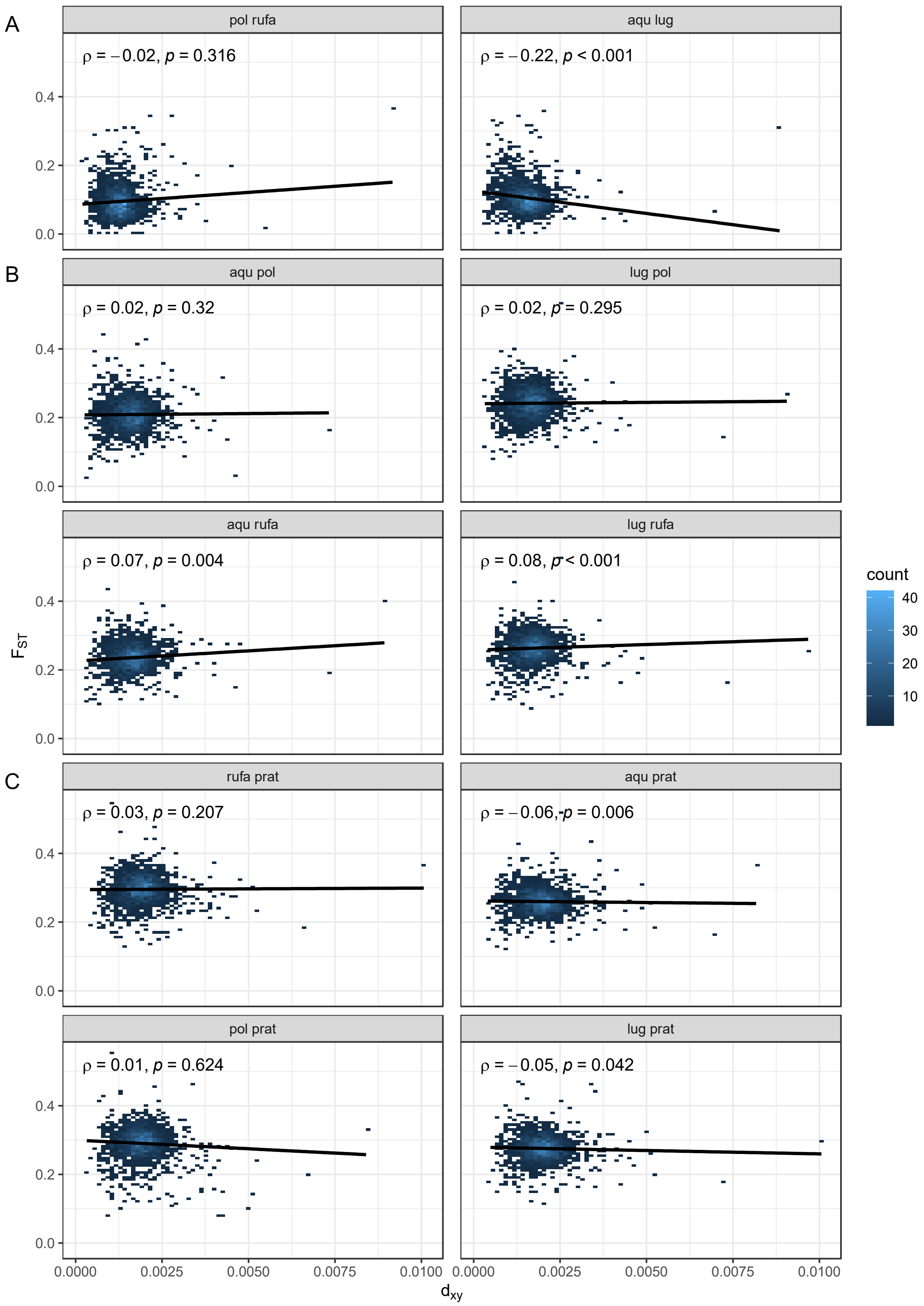


**Fig S4. Correlations between divergence ($d_{xy}$) and differentiation ($F_{ST}$).** Stochastic correlation between divergence (x-axis) and differentiation (y-axis) across the *F. rufa* group, computed in 100 kb genomic windows. **A** Sister species pairs, **B** Between-clade species pairs, **C** All species compared to *F. pratensis.* Species are indicated as aqu = *F. aquilonia*, lug = *F. lugubris*, rufa = *F. rufa*, pol = *F. polyctena*, prat = *F. pratensis.* Correlation tests are performed with Spearman's correlation. A linear regression line with its 95% confidence interval (shading) is shown.
