## Supplementary material for "Introgression and divergence in a young species group": S5 Fig

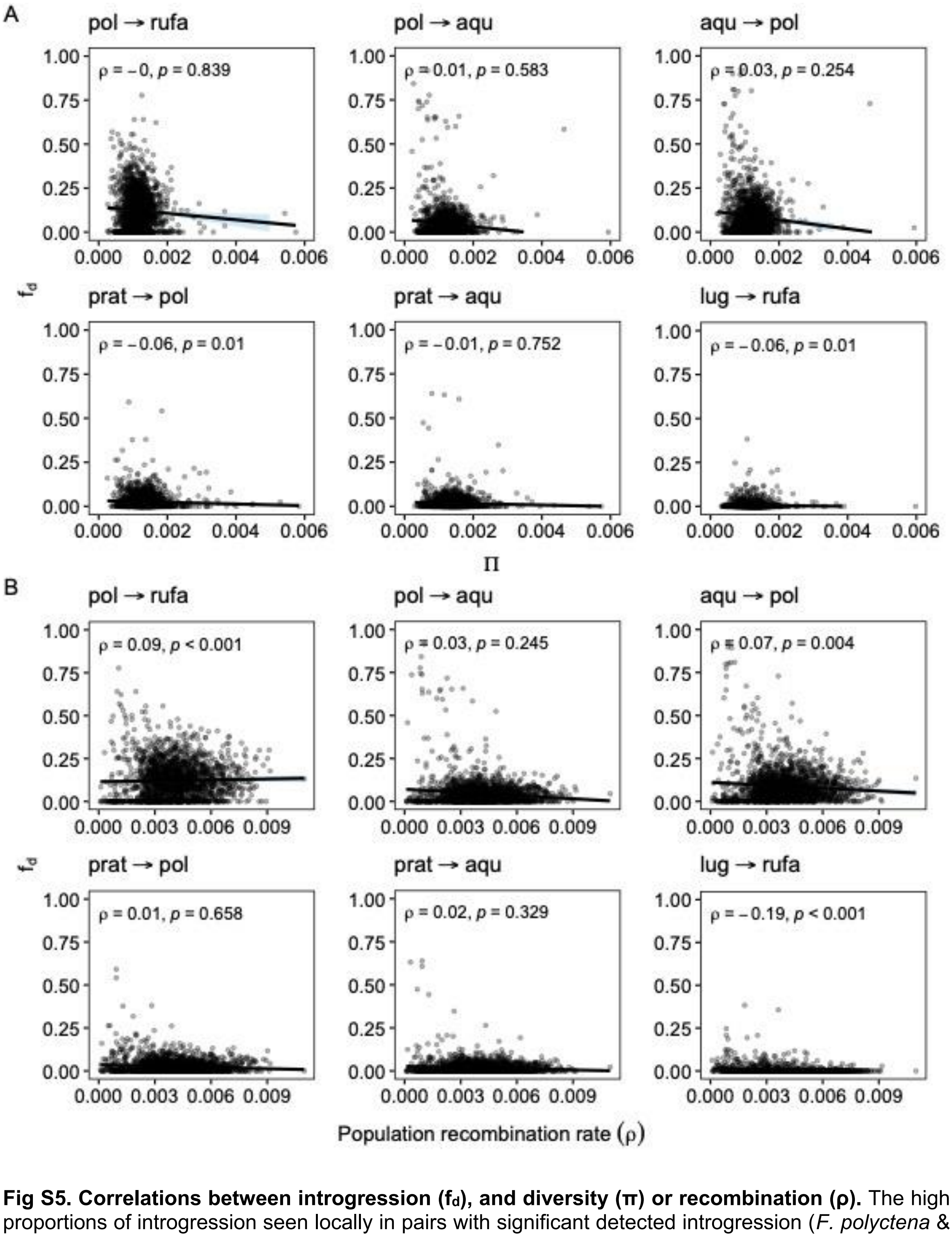


**Fig S5. Correlations between introgression ($f_d$), and diversity (π) or recombination (ρ).** The high proportions of introgression seen locally in pairs with significant detected introgression (*F. polyctena* & *F. rufa*; *F. aquilonia* & *F. polyctena*, bidirectional) are not seen to a similar extent in control pairs (*F. pratensis* & *F. polyctena*; *F. pratensis* & *F. aquilonia*; *F. lugubris* & *F. rufa*). This indicates that the high $f_d$ values are not fully explained by noise due to low information in regions of low effective population size. Values are computed in 100 kb genomic windows. **A** Correlation between diversity (average of the two species) and introgression, **B** Correlation between population recombination rate (average of *F. aquilonia* and *F. polyctena*) and introgression. Species are indicated as aqu = *F. aquilonia*, lug = *F. lugubris*, rufa = *F. rufa*, pol = *F. polyctena*, prat = *F. pratensis.* Correlation tests are performed with Spearman's correlation. A linear regression line with its 95% confidence interval (shading) is shown.
