## Supplementary material for "Introgression and divergence in a young species group": S6 Fig

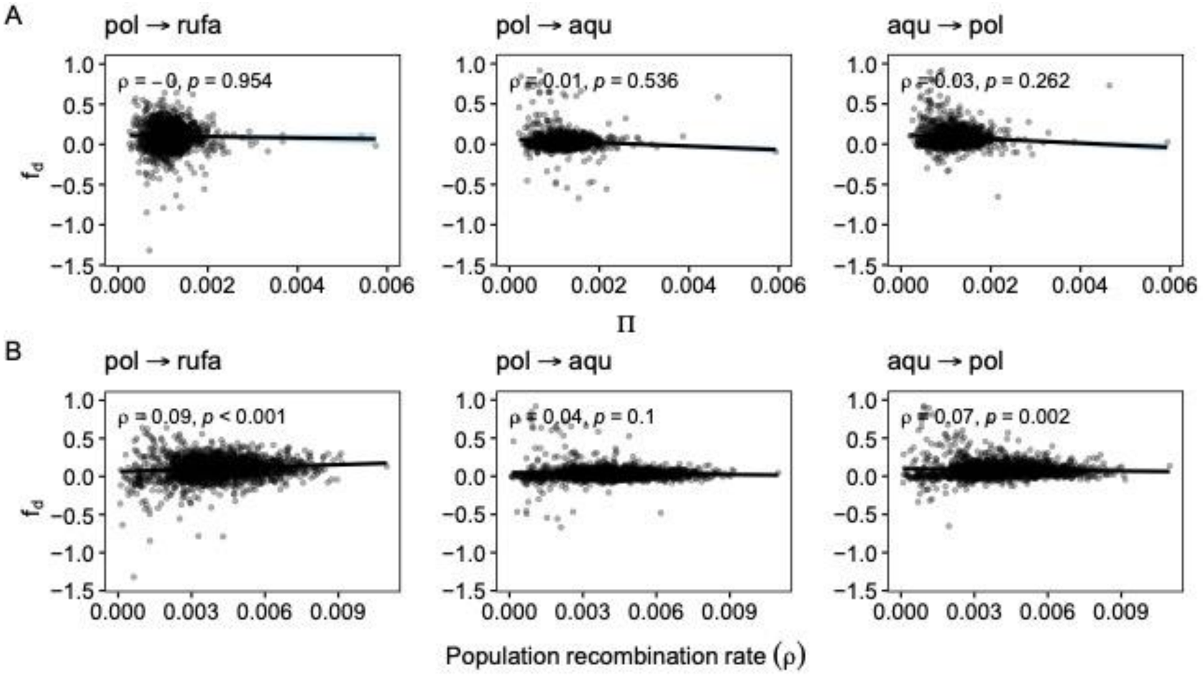


**Fig S6. Correlations between introgression ($f_d$), and diversity (π) or recombination (ρ) with negative $f_d$ values included.** The higher number of not only positive but also negative $f_d$ outliers (a 'funnel-shaped' data distribution) in some comparisons may be indicative of high $f_d$ values caused by increased noise. This noise would arise due to higher drift in regions of low effective population size ($N_e$), as both π and ρ are affected by $N_e$. Species pairs with significant detected introgression are presented. Values are computed in 100 kb genomic windows. **A** Correlation between diversity (average of the two species) and $f_d$, **B** Correlation between population recombination rate (average of *F. aquilonia* and *F. polyctena*) and $f_d$. Species are indicated as aqu = *F. aquiloni*a, rufa = *F. rufa*, pol = *F. polyctena.* Correlation tests are performed with Spearman's correlation. A linear regression line with its 95% confidence interval (shading) is shown.
