## Supplementary material for "Introgression and divergence in a young species group": S1 Table

**Table S1. Detailed information on sampling.** The samples or populations used for each analysis ('x' or a colored cell indicates that the sample was used in the analysis in question). 'Admixed 1' are admixed individuals between *F. aquilonia* and the *F. polyctena/F. rufa* clade. 'Admixed 2' are admixed *F. lugubris* individuals. The species abbreviations in VCF ID column are based on tentative morphology and can be ignored.

| ID | Genomic assignment | Sampling year | Geographic location | Latitude | Longitude | Population | Literature reference for genomic data | VCF ID | NJ-tree | Pi, Dxy, Fst, fd | *f*-branch split | *f*-branch simple | Population recombination rate (computed in ref. 3) | NJ-network |
|---|---|---|---|---|---|---|---|---|---|---|---|---|---|---|
| 44 | *F. aquilonia* | 2006 | Finland | 65.532 | 27.768 | Rantala | 5 | 44-FaquH | X | aqu | aqu_fi | aqu | | X |
| 104 | *F. aquilonia* | 2019 | Finland | 60.580 | 24.037 | Pusula | 5 | 104-Faqu | X | aqu | aqu_fi | aqu | | X |
| 109 | *F. aquilonia* | 2016 | Finland | 69.761 | 26.991 | Kevo | 5 | 109-Faqu | X | aqu | aqu_fi | aqu | | X |
| 106 | *F. aquilonia* | 2019 | Finland | 60.038 | 23.379 | Grabbskog | 5 | 106-Faqu | X | aqu | | aqu | | X |
| 107 | *F. aquilonia* | 2019 | Finland | 60.039 | 23.047 | Solböle | 5 | 107-Faqu | X | aqu | | aqu | | X |
| 108 | *F. lugubris* | 2019 | Finland | 67.526 | 24.981 | Särestöniemi | 5 | 108-Flug | X | lug | lug_fi_1 | lug | | X |
| 113 | *F. lugubris* | 2016 | Finland | 64.555 | 29.907 | Vartius | 5 | 113-Flug | X | lug | lug_fi_1 | lug | | X |
| 11 | *F. lugubris* | 2005 | Finland | 62.567 | 30.965 | Lamminvaara | 5 | 11-Flug | X | lug | lug_fi_2 | lug | | X |
| 115 | *F. lugubris* | 2016 | Russia | 64.485 | 30.184 | Ozero_Kamennoye | 5 | 115-Flug | X | lug | lug_fi_2 | lug | | X |
| 124 | *F. lugubris* | 2008 | Finland | 62.814 | 30.134 | Mökkikylä_2 | 5 | 124-Flug | X | lug | lug_fi_2 | lug | | X |
| Fexs | Outgroup | NA | Finland | 59.83 | 23.27 | Furuskär | 2;3 | Fexs | X | | Outgroup | Outgroup | | |
| CBCH1 | *F. polyctena* | 2018 | Switzerland | 46.681 | 9.657 | Alvaneu | 4 | CBCH1_1w | X | pol | pol_eswi | pol | pol | X |
| CBCH3 | *F. polyctena* | 2018 | Switzerland | 46.681 | 9.657 | Alvaneu | 4 | CBCH3_1w | X | pol | pol_eswi | pol | pol | X |
| CAGa | *F. polyctena* | 2018 | Switzerland | 46.546 | 6.688 | Chalet_a_Gobet | 4 | CAGa_1w | X | pol | pol_wswi | pol | pol | X |
| NAZa | *F. polyctena* | 2018 | Switzerland | 46.660 | 6.684 | Naz | 4 | NAZa_1w | X | pol | pol_wswi | pol | pol | X |
| VDa | *F. polyctena* | 2018 | Switzerland | 46.577 | 6.628 | Vernand_Dessus | 4 | VDa_1w | X | pol | pol_wswi | pol | pol | X |
| 122 | *F. pratensis* | 2008 | Finland | 62.825 | 30.141 | Mökkikylä | 5 | 122-Fprat | X | prat | prat_fi_1 | prat | | X |
| 123 | *F. pratensis* | 2008 | Finland | 62.681 | 29.687 | Puntarikoski | 5 | 123-Fprat | X | prat | prat_fi_1 | prat | | X |
| 6 | *F. pratensis* | 2005 | Finland | 63.497 | 25.752 | Kinnusranta | 5 | 6-Fprat | X | prat | prat_fi_2 | prat | | X |
| 117 | *F. pratensis* | 2015 | Finland | 60.196 | 25.108 | Vuosaari | 5 | 117-Fprat | X | prat | prat_fi_2 | prat | | X |
| 120 | *F. pratensis* | 2015 | Finland | 60.248 | 25.146 | Länsisalmi | 5 | 120-Fprat | X | prat | prat_fi_2 | prat | | X |
| 26 | *F. rufa* | 2005 | Finland | 60.925 | 24.457 | Katiskoski | 5 | 26-Frufa | X | rufa | rufa_fi_1 | rufa | | X |
| 72 | *F. rufa* | 2006 | Finland | 63.360 | 24.358 | Länttäpatti | 5 | 72-Frufa | X | rufa | rufa_fi_1 | rufa | | X |
| 102 | *F. rufa* | 2005 | Finland | 60.680 | 23.593 | Kalliola | 5 | 102-Frufa | X | rufa | rufa_fi_1 | rufa | | X |
| 16 | *F. rufa* | 2005 | Finland | 60.861 | 22.246 | Lännenkulma | 5 | 16-Frufa | X | rufa | rufa_fi_2 | rufa | | X |
| 37 | *F. rufa* | 2005 | Finland | 61.377 | 27.558 | Tyrynmäki | 5 | 37-Frufa | X | rufa | rufa_fi_2 | rufa | | X |
| CBAQ1 | *F. aquilonia* | 2018 | Switzerland | 46.661 | 10.230 | Stabelchod | 4 | CBAQ1_1w | X | | aqu_swi_ | | aqu | X |
| Lai1 | *F. aquilonia* | 2018 | Scotland | 58.028 | 4.441 | Lairg | 4 | Lai_1w | X | | aqu_swi_ | | aqu | X |
| Loa | *F. aquilonia* | 2018 | Scotland | 57.910 | 5.082 | Loch_Achall | 4 | Loa_1w | X | | aqu_swi_ | | aqu | X |
| CBAQ2 | *F. aquilonia* | 2018 | Switzerland | 46.653 | 10.189 | Alp_La_Schera | 4 | CBAQ2_2w | X | | | | aqu | X |
| CBAQ3 | *F. aquilonia* | 2018 | Switzerland | 46.661 | 10.230 | Stabelchod | 4 | CBAQ3_1w | X | | | | aqu | X |
| Lai2 | *F. aquilonia* | 2018 | Scotland | 58.028 | 4.441 | Lairg | 4 | Lai_2w | X | | | | aqu | X |
| 14 | Admixed 1 | 2005 | Finland | 59.921 | 23.040 | Grundsund | 5 | 14-Fpol | | | | | | X |
| 36 | Admixed 1 | 2005 | Finland | 61.256 | 28.674 | Savilahti | 5 | 36-FpolH | | | | | | X |
| 43 | Admixed 1 | 2005 | Finland | 60.703 | 24.689 | Hiivola | 5 | 43-Fpol | | | | | | X |
| 52 | Admixed 1 | 2005 | Finland | 62.565 | 22.794 | Nyrhisperä | 5 | 52-Fpol | | | | | | X |
| 53 | Admixed 1 | 2006 | Finland | 62.309 | 23.651 | Kummunmäki | 5 | 53-Fpol | | | | | | X |
| 55 | Admixed 1 | 2007 | Finland | 62.026 | 26.906 | Luusjoki | 5 | 55-FaquH | | | | | | X |
| 59 | Admixed 1 | 2006 | Finland | 60.838 | 21.808 | Kiili | 5 | 59-FpolH | | | | | | X |
| 65 | Admixed 1 | 2005 | Finland | 60.518 | 27.338 | Rakila | 5 | 65-Frufa | | | | | | X |
| Att1 | Admixed 1 | 2018 | Finland | 60.215 | 19.907 | Åttböle | 4 | Att1_1w | | | | | | X |
| Fis2 | Admixed 1 | 2018 | Finland | 60.151 | 23.557 | Fiskars | 4 | Fis2_1w | | | | | | X |
| Jar6 | Admixed 1 | 2018 | Finland | 60.014 | 20.001 | Järsö | 4 | Jar6_1w | | | | | | X |
| Lok3 | Admixed 1 | 2018 | Finland | 60.375 | 19.810 | Lökholm | 4 | Lok3_1w | | | | | | X |
| RN423 | Admixed 1 | 2022 | Germany, East | 51.411 | 14.518 | Bärwalde | 1 | RN423 | | | | | | X |
| 119 | Admixed 2 | 2010 | Finland | 60.197 | 25.172 | Uutela | 5 | 119-Flug | | | | | | X |
| 125 | Admixed 2 | 2008 | Finland | 62.673 | 29.583 | Härkinvaara | 5 | 125-Flug | | | | | | X |
| 126 | Admixed 2 | 2008 | Finland | 62.746 | 30.037 | Pilkkasuo | 5 | 126-Flug | | | | | | X |
| RN415 | Admixed 2 | 2010 | Switzerland, East | 46.531 | 9.646 | Alp_Flix_1 | 1 | RN415 | | | | | | X |
| RN416 | Admixed 2 | 2010 | Switzerland, East | 46.517 | 9.634 | Alp_Flix_2 | 1 | RN416 | | | | | | X |
| 9 | *F. aquilonia* | 2005 | Finland | 60.505 | 26.233 | Paavalinkylä | 5 | 9-Faqu | X | | | | | X |
| 10 | *F. aquilonia* | 2006 | Finland | 62.608 | 24.971 | Saramäki | 5 | 10-Fpol | X | | | | | X |
| 13 | *F. aquilonia* | 2005 | Finland | 64.517 | 25.123 | Juomala | 5 | 13-Faqu | | | | | | X |
| 17 | *F. aquilonia* | 2005 | Finland | 61.265 | 21.513 | Järvenkulma | 5 | 17-Faqu | X | | | | | X |
| 22 | *F. aquilonia* | 2005 | Finland | 68.064 | 23.484 | Karila | 5 | 22-Faqu | X | | | | | X |
| 24 | *F. aquilonia* | 2006 | Finland | 68.097 | 27.767 | Vuotso | 5 | 24-Faqu | X | | | | | X |
| 25 | *F. aquilonia* | 2005 | Finland | 60.046 | 23.663 | Karjaa | 5 | 25-Faqu | X | | | | | X |
| 27 | *F. aquilonia* | 2005 | Finland | 66.650 | 29.106 | Kaarti | 5 | 27-Fpol | X | | | | | X |

### (Table S1. continues)

| | | | | | | | | | | | | | |
|---|---|---|---|---|---|---|---|---|---|---|---|---|---|
| 30 | *F. aquilonia* | 2008 | Finland | 66.290 | 29.527 | Käylä | 5 | 30-Faqu | X | | | | X |
| 31 | *F. aquilonia* | 2006 | Finland | 67.360 | 29.640 | Salla | 5 | 31-Faqu | X | | | | X |
| 32 | *F. aquilonia* | 2005 | Finland | 62.131 | 27.342 | Väisälä | 5 | 32-Frufa | X | | | | X |
| 38 | *F. aquilonia* | 2005 | Finland | 66.476 | 28.628 | Maaninkavaara | 5 | 38-Fpol | X | | | | X |
| 42 | *F. aquilonia* | 2006 | Finland | 61.621 | 25.311 | Kissamäki | 5 | 42-Faqu | X | | | | X |
| 47 | *F. aquilonia* | 2006 | Finland | 67.165 | 23.581 | Lappea | 5 | 47-FaquH | X | | | | X |
| 48 | *F. aquilonia* | 2005 | Finland | 62.945 | 31.293 | Hattuvaara | 5 | 48-Fpol | X | | | | X |
| 50 | *F. aquilonia* | 2006 | Finland | 60.936 | 25.124 | Saari | 5 | 50-FaquH | X | | | | X |
| 51 | *F. aquilonia* | 2006 | Finland | 60.907 | 24.011 | Ojajärvi | 5 | 51-FaquH | X | | | | X |
| 58 | *F. aquilonia* | 2005 | Finland | 61.604 | 25.054 | Järvenpää | 5 | 58-Fpol | X | | | | X |
| 62 | *F. aquilonia* | 2006 | Finland | 65.945 | 27.708 | Kilsikorpi | 5 | 62-FaquH | X | | | | X |
| 66 | *F. aquilonia* | 2005 | Finland | 65.658 | 27.772 | Karsikkojärvi | 5 | 66-Fpol | | | | | X |
| 67 | *F. aquilonia* | 2005 | Finland | 61.249 | 29.121 | Purnujärvi | 5 | 67-Faqu | X | | | | X |
| 70 | *F. aquilonia* | 2005 | Finland | 64.131 | 30.049 | Elsinmäki | 5 | 70-Faqu | X | | | | X |
| 76 | *F. aquilonia* | 2006 | Finland | 62.084 | 23.436 | Vähä_Mustajärvi | 5 | 76-FaquH | X | | | | X |
| 77 | *F. aquilonia* | 2005 | Finland | 63.633 | 28.570 | Luokkimäki | 5 | 77-Fpol | X | | | | X |
| 83 | *F. aquilonia* | 2006 | Finland | 63.104 | 24.105 | Latvalampi | 5 | 83-Faqu | X | | | | X |
| 91 | *F. aquilonia* | 2006 | Finland | 63.728 | 29.716 | Kataja | 5 | 91-FpolH | X | | | | X |
| 93 | *F. aquilonia* | 2005 | Finland | 67.213 | 26.848 | Joutsela | 5 | 93-Faqu | | | | | X |
| 99 | *F. aquilonia* | 2005 | Finland | 63.376 | 26.895 | Lampaanjärvi | 5 | 99-Faqu | X | | | | X |
| 103 | *F. aquilonia* | 2005 | Finland | 63.119 | 29.393 | Pihlajavaara | 5 | 103-Faqu | X | | | | X |
| 114 | *F. aquilonia* | 2016 | Finland | 64.548 | 29.896 | Vartius_2 | 5 | 114-FaquH | X | | | | X |
| CF14a | *F. aquilonia* | 2018 | Finland | 62.635 | 26.201 | Pukara | 4 | CF14a_1w | X | | | | X |
| CF4b | *F. aquilonia* | 2018 | Finland | 63.503 | 26.087 | Koivula | 4 | CF4b_1w | X | | | | X |
| CF8b | *F. aquilonia* | 2018 | Finland | 63.682 | 27.545 | Sonkajärvi | 4 | CF8b_1w | X | | | | X |
| Pus2 | *F. aquilonia* | 2018 | Finland | 60.581 | 24.038 | Pusula | 4 | Pus2_1w | X | | | | X |
| 1 | *F. lugubris* | 2006 | Finland | 61.602 | 21.907 | Noormarkku | 5 | 1-Fprat | X | | | | X |
| 56 | *F. lugubris* | 2006 | Finland | 66.662 | 27.735 | Soppela | 5 | 56-Flug | X | | | | X |
| 88 | *F. lugubris* | 2005 | Finland | 61.207 | 23.089 | Anttilankulma | 5 | 88-Flug | X | | | | X |
| CBCH2 | *F. polyctena* | 2018 | Switzerland | 46.681 | 9.657 | Alvaneu | 4 | CBCH2_2w | X | | | | X |
| 2 | *F. pratensis* | 2005 | Finland | 62.005 | 26.080 | Kaakkomäki | 5 | 2-Fprat | X | | | | X |
| 3 | *F. pratensis* | 2006 | Finland | 62.283 | 22.046 | Järviharju | 5 | 3-Fprat | X | | | | X |
| 12 | *F. rufa* | 2006 | Finland | 61.772 | 23.709 | Leppälahti | 5 | 12-Frufa | | | | | X |
| RN419 | *F. rufa* | 2022 | Germany, West | 50.464 | 8.209 | Gaudernbach | 1 | RN419 | X | | | | X |
| RN420 | *F. rufa* | 2022 | Germany, West | 50.420 | 8.332 | Weilmünster | 1 | RN420 | | | | | X |
| RN424 | *F. rufa* | 2022 | Germany, East | 51.254 | 14.781 | Diehsa | 1 | RN424 | | | | | X |


**Literature reference for genomic data:**

1 This study

2 Dhaygude K, Nair A, Johansson H, Wurm Y, Sundström L. (2019).
The first draft genomes of the ant Formica exsecta, and its Wolbachia endosymbiont reveal extensive gene transfer from endosymbiont to host.
BMC Genom. 2019;20:301. pmid:30991952

3 Nouhaud P, Martin SH, Portinha B, Sousa VC, Kulmuni J. (2022).
Rapid and predictable genome evolution across three hybrid ant populations.
PLoS Biol 20(12): e3001914. https://doi.org/10.1371/journal.pbio.3001914

4 Portinha, B., Avril, A., Bernasconi, C., Helantera, H., Monaghan, J., Seifert, B., Sousa, V. C., Kulmuni, J., & Nouhaud, P. (2022).
Whole-genome analysis of multiple wood ant population pairs supports similar speciation histories,
but different degrees of gene flow, across their European ranges.
Molecular Ecology, 31, 3416–3431. https://doi. org/10.1111/mec.16481

5 Satokangas, I., Nouhaud, P., Seifert, B., Punttila, P., Schultz, R., Jones, M. M., Sirén, J., Helanterä, H., & Kulmuni, J. (2023).
Semipermeable species boundaries create opportunities for gene flow and adaptive potential.
Molecular Ecology, 32, 4329–4347. https://doi.org/10.1111/mec.16992
