## Supplementary material for "Introgression and divergence in a young species group": S2 Table

**Table S2. Per-individual coverage.** Average read depth for the newly sequenced individuals computed from bam files after overlap clipping.

| **Individual** | **Coverage** |
|---|---|
| RN415 | 9.32 |
| RN416 | 10.41 |
| RN419 | 12.22 |
| RN420 | 11.30 |
| RN423 | 9.88 |
| RN424 | 9.68 |
