## Supplementary material for "Introgression and divergence in a young species group": S3 Table

**Table S3. Association of low negative $f_d$ values, and diversity (π) or recombination (ρ).** Locally in the genome low population recombination rates are associated with low negative $f_d$ values (significant test statistic). The association of low population recombination rates and both negative and positive outlier $f_d$ values (Table 2) may be caused by increased noise in $f_d$. Such noise can arise due to increased drift in regions of low local effective population size and nonindependence of genomic loci among low recombining regions. The association of diversity and outlier $f_d$ regions cannot, however, be explained by noise alone. The data is computed in genomic windows of 100 kb and tested with chi-squared test, N(aqu pol)=1930. N(pol rufa)=1909. pol = *F. polyctena*, aqu = *F. aquilonia*, rufa = *F. rufa.*
* = significant at 0.05 level, ** = significant at 0.005 level

| Introgressing species and the direction of gene flow | % of low outlier introgression windows ($f_d$ threshold) | Count of the low $f_d$ outlier windows in each bin | | Diversity (π) | | | Population recombination rate (ρ) | | |
|---|---|---|---|---|---|---|---|---|---|
| | | π: low/ high | ρ: low/ high | $X^2$ | df | p-value | $X^2$ | df | p-value |
| pol → aqu | 1% (-0.109) | 9/11 | 18/2 | 0.0495 | 1 | **0.824** | 11.384 | 1 | **< 0.001 **** |
| | 3% (-0.066) | 32/26 | 42/16 | 0.4498 | 1 | **0.503** | 11.137 | 1 | **< 0.001 **** |
| aqu → pol | 1% (-0.107) | 12/8 | 17/3 | 0.4579 | 1 | **0.499** | 8.552 | 1 | **0.003 **** |
| | 3% (-0.045) | 37/21 | 42/15 | 4.0156 | 1 | **0.045 *** | 12.249 | 1 | **< 0.001 **** |
| pol → rufa | 1% (-0.317) | 13/7 | 17/2 | 1.263 | 1 | **0.261** | 10.42 | 1 | **0.001 **** |
| | 3% (-0.151) | 41/17 | 44/13 | 9.4066 | 1 | **0.002 **** | 16.276 | 1 | **< 0.001 **** |
